# Cobalt-sulfur coordination chemistry drives B_12_ loading onto methionine synthase

**DOI:** 10.1101/2023.07.25.550549

**Authors:** Romila Mascarenhas, Arkajit Guha, Zhu Li, Markus Ruetz, Sojin An, Javier Seravalli, Ruma Banerjee

## Abstract

Cobalt-sulfur (Co-S) coordination is labile to both oxidation and reduction chemistry and is rarely seen in Nature. Cobalamin (or vitamin B_12_) is an essential cobalt-containing organometallic cofactor in mammals, and is escorted via an intricate network of chaperones to a single cytoplasmic target, methionine synthase. In this study, we report that the human cobalamin trafficking protein, MMADHC, exploits the chemical lability of Co-S coordination, for cofactor off-loading onto methionine synthase. Cys-261 on MMADHC serves as the β-axial ligand to cobalamin. Complex formation between MMADHC and methionine synthase is signaled by loss of the lower axial nitrogen ligand, leading to five-coordinate thiolato-cobalamin. Nucleophilic displacement by the vicinal thiolate, Cys-262, completes cofactor transfer to methionine synthase and release of a cysteine disulfide-containing MMADHC. The physiological relevance of this mechanism is supported by clinical variants of MMADHC, which impair cofactor binding and off-loading, explaining the molecular basis of the associated homocystinuria.

---

Metal redox states and coordination chemistry are tightly regulated in trafficking pathways for moving metal and organometallic cargo to target proteins. Vitamin B_12_ or cobalamin is a large organometallic cofactor that relies on a host of chaperones to tailor and deliver it to two B_12_-dependent enzymes in humans: methylcobalamin (MeCbl)-dependent cytoplasmic methionine synthase (MTR) and 5’-deoxyadenosylcobalamin (AdoCbl)-dependent mitochondrial methylmalonyl-CoA mutase (MMUT)^1^. Inherited defects in B_12_ trafficking lead to combined or isolated homocystinuria and methylmalonic aciduria, depending on whether the affected locus is in the common or branched segments in the pathway^2^. Studies spanning a decade have provided detailed structural and mechanistic insights into the strategies deployed by the mitochondrial chaperones for cofactor loading to and off-loading from MMUT^3, 4^. In contrast, much less is known about the mechanism of B_12_ delivery to MTR and the role of the MMADHC (or CblD) chaperone.

MTR catalyzes two successive methyl transfer reactions from MeCbl to homocysteine and from N^5^-methytetrahydrofolate (CH_3_-H_4_F) to cob(I)alamin, generating methionine and tetrahydrofolate (H_4_F), respectively (Figure 1A). In this process, MTR converts CH_3_-H_4_F, the circulating form of folic acid, to H_4_F, making it available for purine and amino acid biosynthesis reactions^5^. The cofactor cycles between the MeCbl and supernucleophilic cob(I)alamin forms, which is susceptible to inactivation once every ∼2000 turnovers^6^. Inactive MTR is repaired via a reductive methylation reaction, which uses S-adenosylmethionine (AdoMet) as a methyl donor and NADPH-dependent methionine synthase reductase (MTRR) a diflavin oxidoreductase, as a dedicated redox partner^7^. While MTRR has been postulated to assist in cofactor loading onto MTR^8^, the physiological relevance of this observation is uncertain^9^.

**Figure 1.**
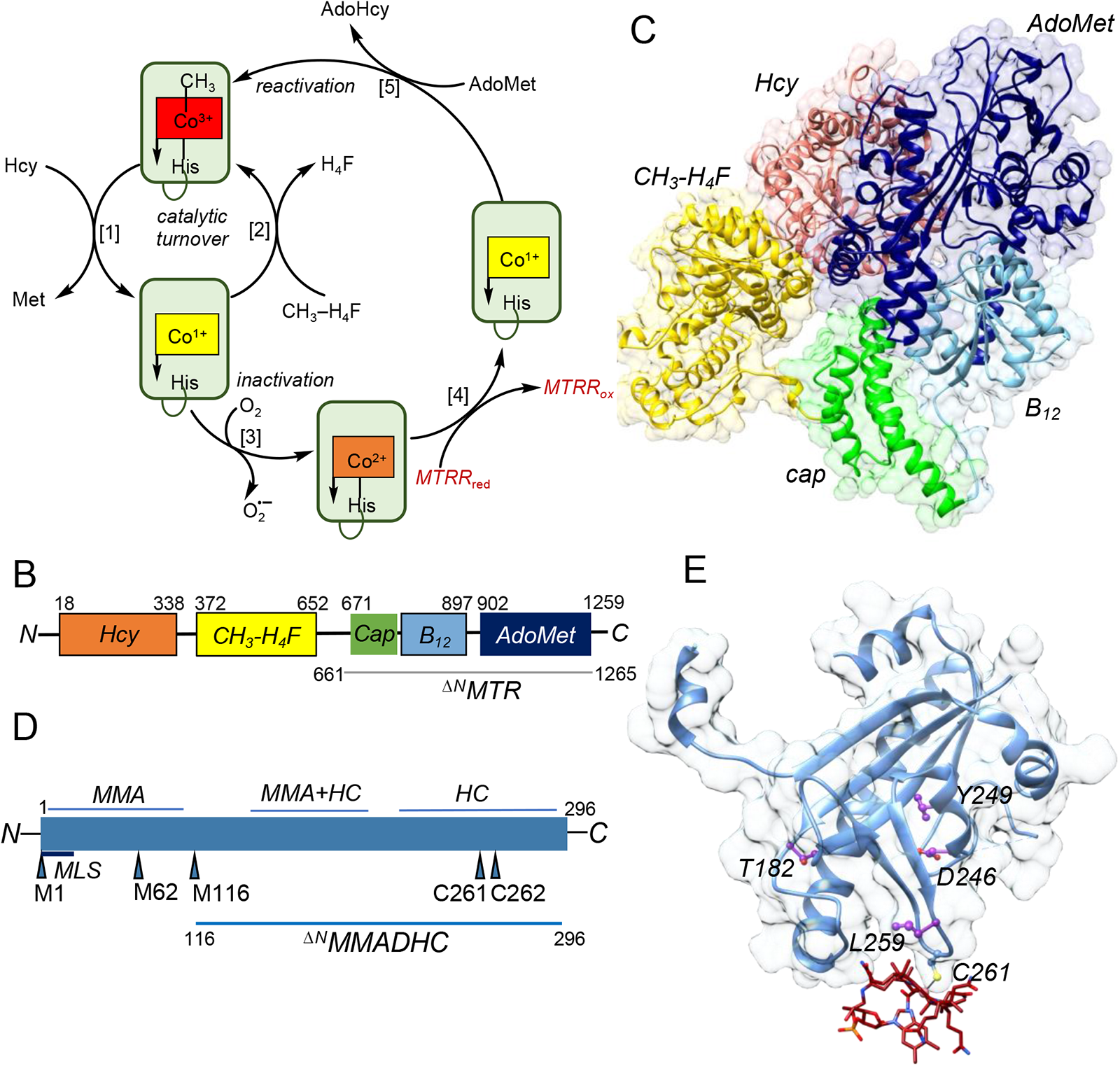
Structures and reaction mechanism of MTR and MMADHC. **A**. MTR catalyzes two methyl transfer reactions from MeCbl to homocysteine (Hcy) to produce methionine [*step* 1], and from CH_3_-H_4_F to cob(I)alamin to produce H_4_F [2]. When cob(I)alamin is accidentally oxidized to cob(II)alamin [3], inactive MTR is repaired by NADPH-dependent MTRR [4] and AdoMet [5]. **B.** Cartoon representing the modular organization of MTR. **C**. ^ΔN^MTR used in this study, comprises the cap, B_12_ and AdoMet domains (residues 661-1265) but lacks the homocysteine (Hcy) and CH_3_-H_4_F domains. **C**. AlphaFold model of human MTR is shown as ribbons and surface view in the same color scheme as in B. The model represents the reactivation conformation in which the cap domain is adjacent to the B_12_-domain and the activation domain is positioned above the B_12_-domain. **D.** Cartoon representation of MMADHC. Mutations in the N-and C-terminal thirds of the protein are associated with isolated methylmalonic aciduria (MMA) and homocystinuria (HC), respectively while mutations in the middle are associated with both diseases. **E**. Structure of ^ΔN^MMADHC-thiolato-cob(III)alamin (PDB 6X8Z) shown in ribbon and surface view. Cys-261 provides the sulfur ligand to B_12_ (red sticks). Mutations that lead to HC are shown as purple ball and sticks.

Human MTR is an ∼140 KDa monomeric protein, which is predicted to be modular in organization like the *E.coli* homolog (Figure 1B). A recent structural characterization of *E.coli* MTR revealed that in its resting state, the B_12_ domain is nestled between the CH_3_–H_4_F and homocysteine domains, while the AdoMet domain is conformationally dynamic^10^. In the reactivation cycle, the AdoMet domain engages with the B_12_ domain as seen in crystal structures of the C-terminal half of the protein^11^. The AlphaFold model of full-length MTR represents the reactivation conformation in which the AdoMet domain protects the B_12_-binding site from solvent exposure while the cap domain is disengaged^19^ (Figure 1C). Studies on human MTR have been thwarted by the inability to express the full-length protein in sufficient quantities for biochemical studies. Furthermore, the mechanism by which B_12_ is loaded onto MTR, and the roles of the cytoplasmic chaperones MMACHC and MMADHC, which are absent in bacteria, are unknown.

The MMACHC (or CblC) chaperone is most frequently mutated in patients with inborn errors of cobalamin metabolism^12^. MMACHC has broad substrate and reaction specificity and catalyzes β-ligand elimination via the glutathione-dependent dealkylation of alkyl-cobalamins^13^, denitration of nitrocobalamin^14^ or reductive elimination of cyano-(CNCbl)^15^ or aquo-cobalamin (H_2_OCbl)^16^. The resulting cob(II)alamin product is stabilized on MMACHC and forms an interprotein cobalt-sulfur (Co-S) coordination complex with MMADHC^17^. The structure of MMADHC lacking the first 133 amino acids (^ΔN^MMADHC), which are predicted to be disordered, most closely resembles MMACHC^17, 18^, which also belongs to the flavin nitroreductase superfamily.

MMADHC variants represent the most complex class of inherited defects in B_12_ metabolism^19^. Mutations in the N-and C-terminal thirds of the protein are associated with isolated methylmalonic aciduria and homocystinuria aciduria, respectively while mutations in the middle are associated with both diseases (Figure 1D)^20, 21^. MMADHC harbors two internal translation initiation sites at Met-62 and Met-116 and variants lacking the N-terminal 115 amino acids are still capable of supporting MTR^20^. While MMADHC was initially reported to be incapable of binding B_12_ ^22, 23^, the fully reduced protein was later shown to coordinate cobalamin via an unusual cobalt-sulfur (Co-S) bond involving Cys-261^17^ (Figure 1E).

The physiological relevance of the Co-S bond, which is relatively rare in Nature, and its role in B_12_ trafficking was previously unknown. Herein, we report that the Co-S bond in the MMADHC-cobalamin complex is instrumental for loading the cofactor onto MTR. We furnish evidence in support of a cobalamin loading mechanism in which the Cys-262 thiolate attacks the Co-S-Cys^261^ bond in the MMADHC-S-cob(III)-MTR complex, forming a disulfide-containing MMADHC, and displacing cofactor-loaded MTR. We demonstrate that a subset of clinical MMADHC variants impact the kinetics of Co-S coordination and formation of the MTR-MMADHC complex, revealing the mechanistic underpinnings of disease.

## Results

We focused our studies on the C-terminal half of MTR, comprising the B_12_ and AdoMet-binding domains (denoted hereon as ^ΔN^MTR, Figure 1B), due to the low soluble expression yield of recombinant human MTR in *E.coli* or insect cells. Addition of ^ΔN^MTR to an anaerobic solution of cob(II)alamin elicited a blue shift from 474 nm to 470 nm (Figure S1A). ^ΔN^MTR also bound MeCbl from solution but not OH_2_Cbl or glutathionyl cobalamin (GSCbl) (Figure S1B-D).

To test whether ^ΔN^MTR is sufficient to support reductive methylation, we mixed ^ΔN^MTR • cob(II)alamin with MTRR, NADPH and AdoMet (Figure 1A). HPLC analysis of the reaction mixture confirmed formation of MeCbl (Figure S1E). ^ΔN^MTR-bound cob(II)alamin was characterized by electron paramagnetic resonance (EPR) spectroscopy, which affords ready distinction between the presence of an axial nitrogen versus an oxygen ligand. Interactions between the unpaired electron and the cobalt nucleus (I=^7^/_2_) results in hyperfine splitting into an 8-line spectrum while nitrogen (I=1) coordination leads to superhyperfine splitting of each of the eight lines into triplets. The triplet structures are clearly visible in the high-field region of the spectrum (3250-4000 G) of free cob(II)alamin where the dimethylbenzimidazole (DMB) nitrogen serves as the lower axial ligand (Figure S2A). In the presence of ^ΔN^MTR, the EPR spectrum comprises a mixture of 5-coordinate cob(II)alamin with an axial oxygen (∼55%) or nitrogen (45%) ligand (Figure S2B,E). Cobalamins bind to MTR in a “base-off” conformation (where the base refers specifically to the DMB ligand), and the lower axial coordination site can be occupied by His-785 (Figure S2G). The EPR spectrum indicates that cob(II)alamin bound to ^ΔN^MTR exists as an equilibrium mixture containing a histidine or a water ligand. Introduction of the H785A mutation in ^ΔN^MTR, confirmed this assignment by revealing a single population of cob(II)alamin, i.e. with an oxygen ligand, resembling the spectrum of “base-off” cob(II)alamin in solution (Figure S2C,D,F).

Next, we tested whether MMACHC or MMADHC can load cobalamin onto MTR. However, mixing ^ΔN^MTR with MMACHC loaded with either cob(II)alamin or MeCbl did not elicit any spectral changes (data not shown), and analytical gel filtration chromatography confirmed retention of cobalamin on MMACHC (Figure S3A). ^ΔN^MMADHC binds OH_2_Cbl from solution to form a thiolato-cob(III)alamin complex (referred to hereon as ^ΔN^MMADHC-S-cob(III)) (Figure 2A), in which either Cys-261 or Cys-262 can provide the sulfur ligand^17^. Mixing ^ΔN^MMADHC-S-cob(III) (λ_max_= 335, 370, 525 and 560 nm) with ^ΔN^MTR resulted in a blue shift with absorption maxima at 360, 495, and 525 nm with isosbestic points at 338, 358, 441 and 525 nm, signaling OH_2_Cbl formation (Figure 2B). The *k_ob_*_s_ (1.0 ± 0.1 min^-1^) for OH_2_Cbl formation was estimated from the fast phase of the biphasic reaction, which accounts for 52% and 82 % of the amplitude change at 350 nm and 555 nm, respectively. Cofactor transfer from ^ΔN^MMADHC-S-cob(III) to ^ΔN^MTR was confirmed by size exclusion chromatography, monitoring the eluant at 280 nm (for protein) and 355 nm (for cobalamin) (Figure 2C). When the same transfer reaction was set up under anaerobic conditions, cob(II)alamin (λ_max_ = 470 nm, isosbestic points at 502 and 388 nm) was stabilized on ^ΔN^MTR (Figure S3B). This represents the first demonstration of B_12_ transfer from the MMADHC chaperone to MTR and its relevance to full-length MTR, was established with the limited amounts of this recombinant protein obtained from insect cell culture^24^. Size exclusion chromatography revealed cobalamin loading onto MTR under aerobic conditions (Figure S3C), confirming the relevance of ^ΔN^MTR as a useful model.

**Figure 2.**
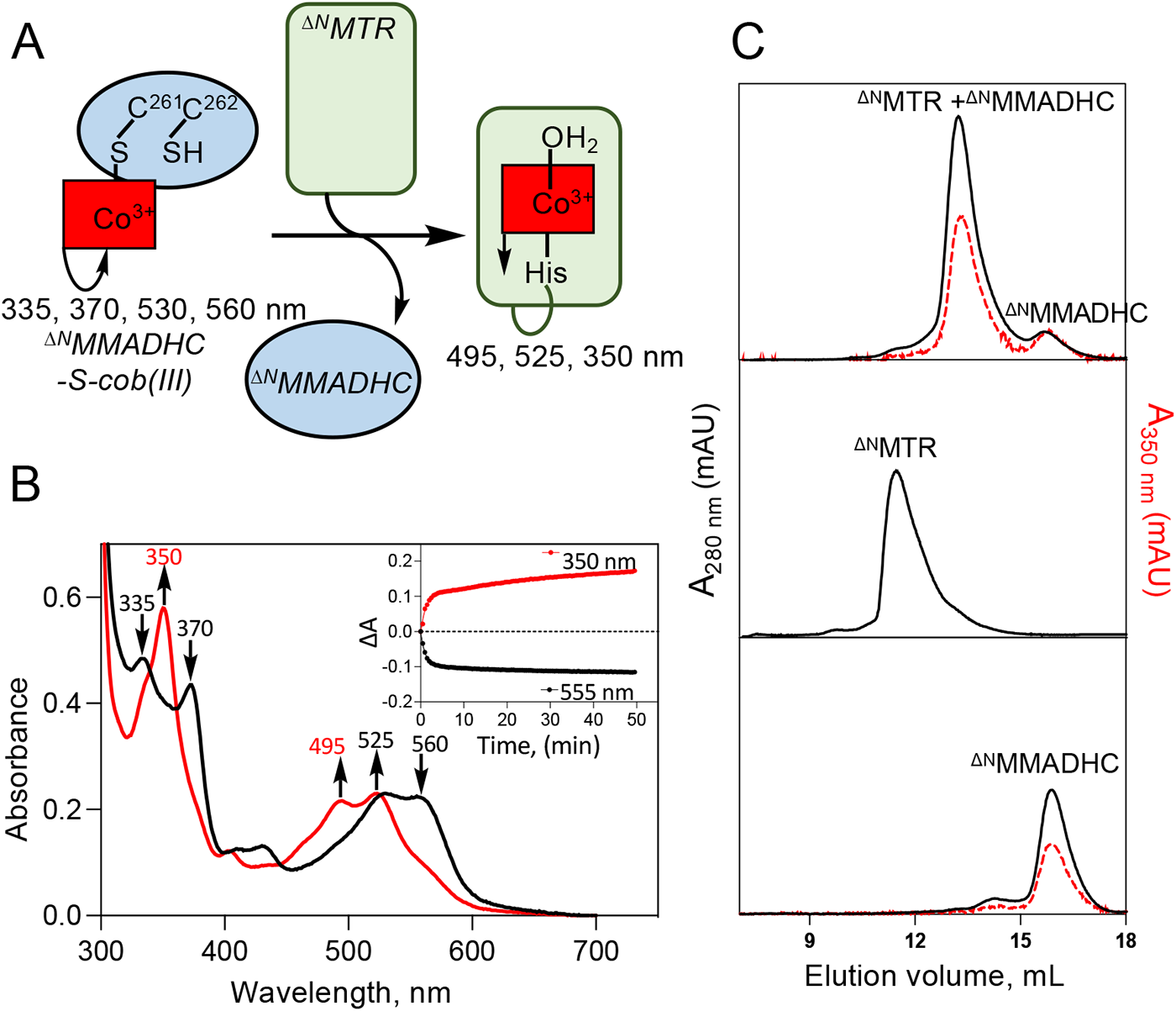
B_12_ transfer from ^ΔN^MMADHC-S-cob(III) to ^ΔN^MTR. **A**. Scheme showing the experimental set-up. **B**. ^ΔN^MMADHC-S-Cob(III) (20 μM, black) was mixed with ^ΔN^MTR (40 μM) in Buffer A at 20°C. Spectra were recorded every 30 s for 50 min and the final (red) spectrum is shown. *Inset.* Time-dependent changes in A_350nm_ (red) and A_555nm_ (black) are shown. **C.** Size exclusion chromatography of the reaction mixture from B and ^ΔN^MTR and ^ΔN^MMADHC-S-cob(III) standards with monitoring at 280 nm (black) and 350 nm (red dashes) confirms the presence of cobalamin bound to ^ΔN^MTR. The data are representative of 3 independent experiments. mAU denotes milli-absorbance units.

We evaluated the roles of Cys-261 and Cys-262 in cofactor transfer from ^ΔN^MMADHC to ^ΔN^MTR. When cofactor-loaded C261S ^ΔN^MMADHC (λ_max_=530, 560, 335 and 370 nm) was mixed with ^ΔN^MTR, the spectrum was virtually unchanged even in the presence of a 3-fold excess of ^ΔN^MTR (Figure 3A). In contrast, when the cofactor-loaded C262S ^ΔN^MMADHC was mixed with ^ΔN^MTR, large spectral changes were observed with maxima at 413 nm and 510 nm and isosbestic points at 389 and 519 nm (Figure 3B). This new spectrum, assigned as base-off thiolato-cob(III)alamin^25^, indicates stabilization of an interprotein ^ΔN^MMADHC-^ΔN^MTR complex via a Co-S bond. A 1:1 stoichiometry for complex formation between C262S ^ΔN^MMADHC and ^ΔN^MTR was observed, with a *K*_D_ estimated to be <50 nM (Figure 3C). Formation of the ^ΔN^MMADHC-^ΔN^MTR complex was confirmed by size exclusion chromatography (Figure 3D).

**Figure 3.**
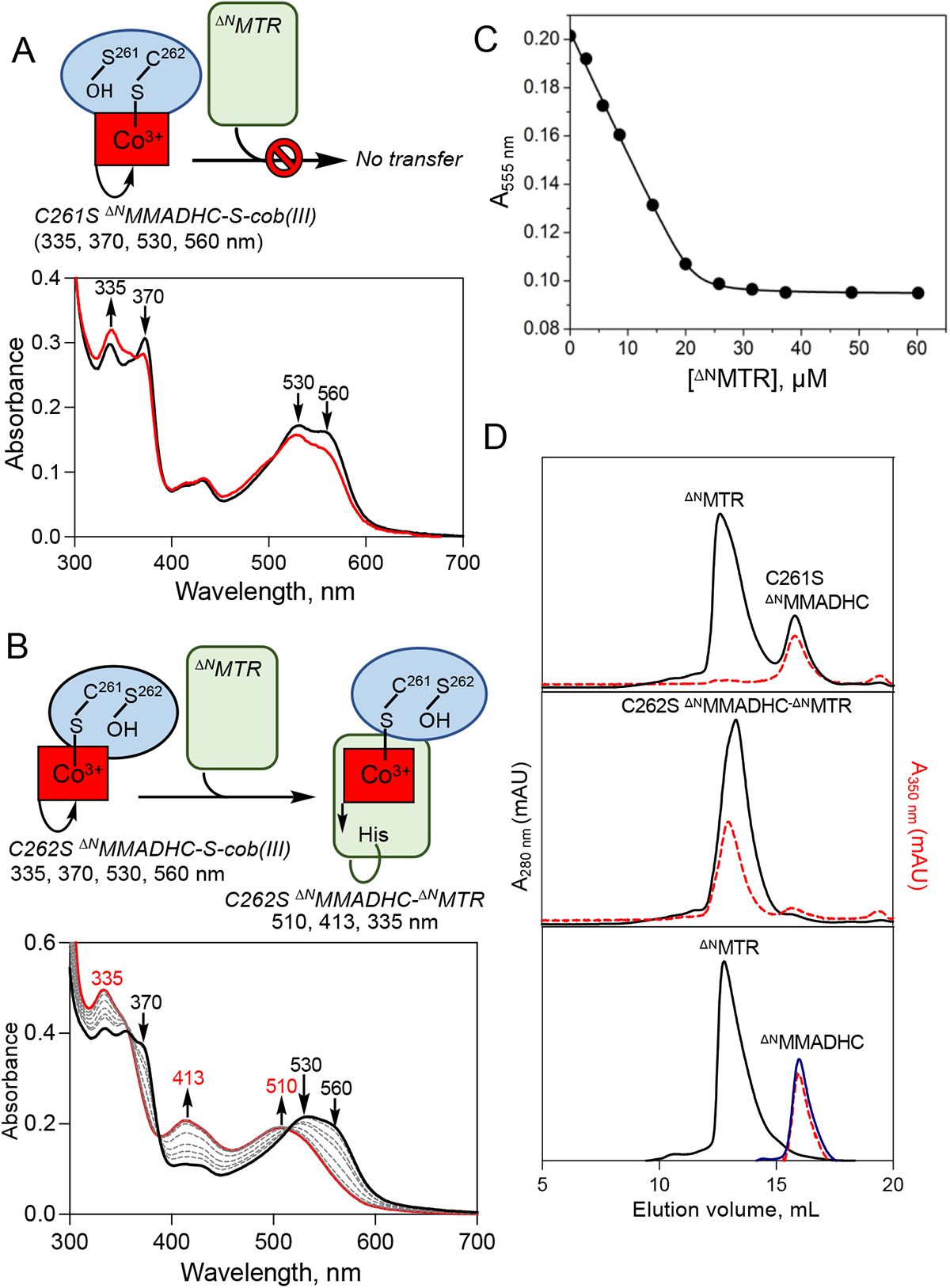
Cofactor transfer from ^ΔN^MMADHC-S-cob(III) to ^ΔN^MTR. **A**. C261S ^ΔN^MMADHC, fails to transfer B_12_ ^ΔN^MTR. Mixing C261S ^ΔN^MMADHC-S-cob(III) (20 μM, black) with 40 μM ^ΔN^MTR in Buffer A at 20°C did not lead to significant spectral changes after 50 min (red trace). **B** C262S ^ΔN^MMADHC-S-cob(III) forms a stable complex with ^ΔN^MTR. Mixing C262S ^ΔN^MMADHC-S-cob(III) (20 μM, black) with 60 μM ^ΔN^MTR at 20°C led to a dramatic spectral shift. The final spectrum at 60 min is in red. The grey lines represent spectra obtained with different concentrations of ^ΔN^MTR (2.5-60 µM). The spectrum shift signals formation of base-off thiolato-cob(III)alamin and formation of a stable interprotein thiolato-cobalamin complex. **C**. The stoichiometry of the inteprotein complex was estimated from the change in A_550nm_ versus concentration of ^ΔN^MTR. **D.** Size exclusion chromatography of the reaction mixtures in A and B, and of ^ΔN^MTR (40 µM) and ^ΔN^MMADHC-S-Cob(III) (20 µM) standards confirmed B_12_ loading onto ^ΔN^MTR from C262S but not C261S ^ΔN^MMADHC. The data are representative of 3 independent experiments.

Our results suggest a mechanism for cofactor docking in which nucleophilic displacement of the Co-S-Cys^261^ bond by the thiolate of Cys-262 leads to elimination of cob(I)alamin on MTR and a disulfide bond between Cys-261 and Cys-262 on MMADHC (Figure 4A). To test this model, we monitored cofactor transfer under anaerobic conditions in the presence of AdoMet, which led to spectral changes (λ_max_= 530 nm) consistent with formation of 6-coordinate MeCbl (Figure 4B). The *k_ob_*_s_ for MeCbl formation (1.2 ± 0.04 min^-1^) was estimated from the fast phase of the biphasic reaction at 570 nm, which is associated with 50% of the amplitude change (Figure 4B, inset). The concentration of MeCbl formed (15 ± 2 µM) was estimated using Δε_570nm_ = 3.5 mM^-1^ cm^-1^, corresponding to a transfer efficiency of 76 ± 4%. HPLC analysis of the reaction products confirmed MeCbl synthesis (Figure 4C). Cofactor transfer from ^ΔN^MMADHC to ^ΔN^MTR was confirmed by analytical gel filtration analysis (Figure S3D).

**Figure 4.**
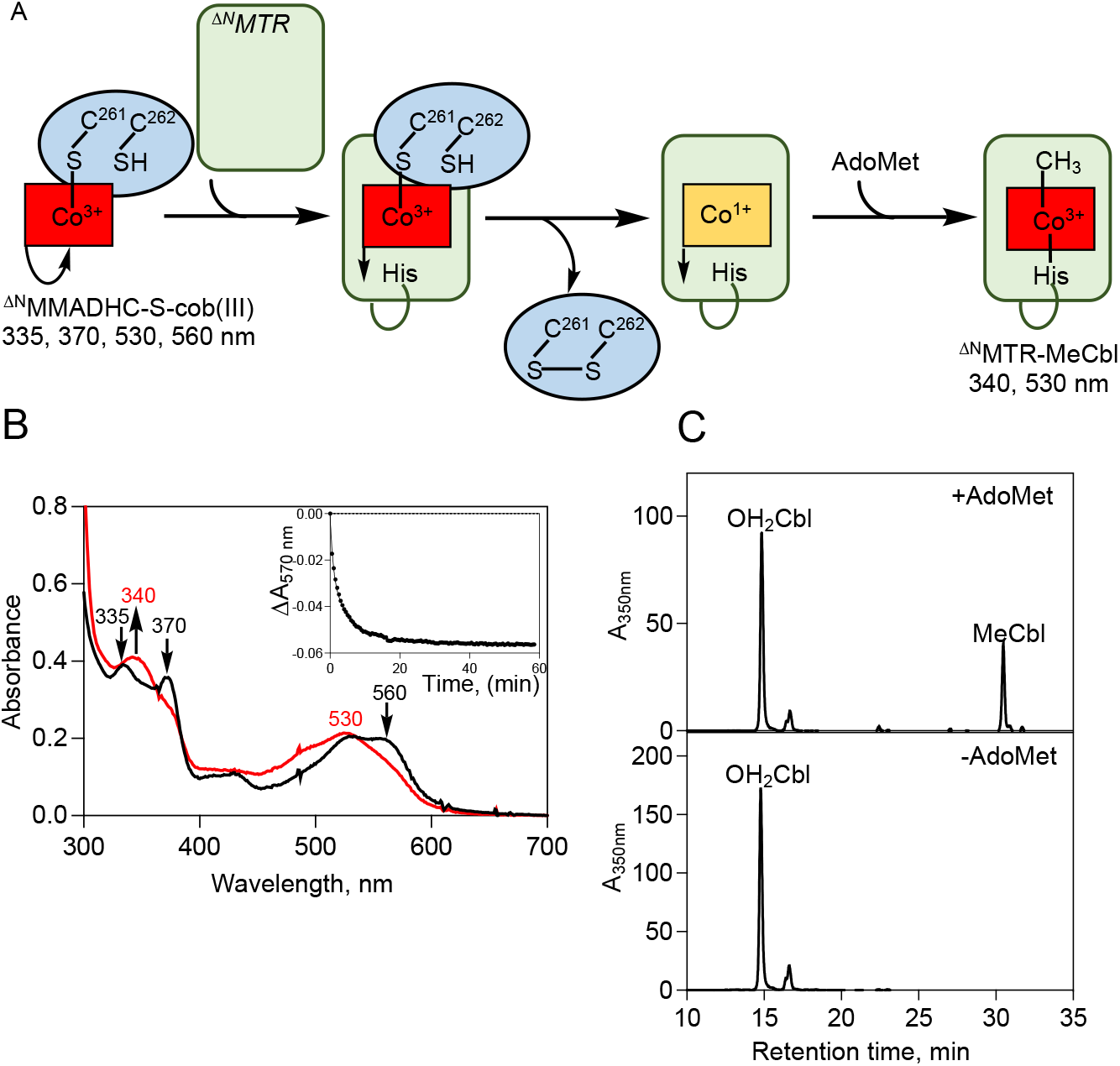
Mechanism of cobalamin transfer from ^ΔN^MMADHC to ^ΔN^MTR. **A**. Postulated mechanism of cobalamin transfer from ^ΔN^MMADHC to ^ΔN^MTR in the presence of AdoMet. **B**. ^ΔN^MMADHC-S-cob(III) (20 μM) and 500 μM AdoMet (black trace) were mixed with ^ΔN^MTR (40 μM) in Buffer A at 20°C under anaerobic conditions. UV/visible spectra were recorded every 30 s for 60 min and the final spectrum (red trace) is shown. *Inset*. Dependence of A_570nm_ on time yielded an estimate of *k_obs_*, the lumped rate constant for cofactor transfer and methylation. **C.** Formation of MeCbl was confirmed by HPLC analysis of the reaction mixture in B. The data are representative of 3 independent experiments.

The role, if any, of His-785, the conserved lower axial ligand to cobalamin on MTR (Figure S2G), in the transfer reaction was investigated next. Mixing cofactor-loaded ^ΔN^MMADHC with H785A ^ΔN^MTR led to a blue shift in the spectrum to 470 nm, indicating cob(II)alamin formation (Figure S4A). The *k_ob_*_s_ (0.99 ± 0.16 min^-1^) for cob(II)alamin formation was estimated from the fast phase of the biphasic reaction at 470 nm or 555 nm, which was associated with 65% of the amplitude change (Figure S4A, *inset*) and is comparable to the value obtained with ^ΔN^MTR. Mixing His-785 ^ΔN^MTR with cofactor-loaded ^ΔN^MMADHC in the presence of AdoMet under anaerobic conditions, resulted in a blue shift in the spectrum (∼460 nm), indicating base-off MeCbl formation (Figure S4B). MeCbl synthesis on His-785 ^ΔN^MTR was confirmed by size exclusion chromatography and HPLC analyses (Figure S4C,D), indicating that His-785 is not involved in B_12_ loading and subsequent MeCbl synthesis on ^ΔN^MTR.

Mass spectrometry (MS) was used to detect formation of the disulfide on ^ΔN^MMADHC between Cys-261 and Cys-262 (Figure 4A). Since disulfides in peptides are difficult to detect and ^ΔN^MMADHC has five cysteines in total, which can scramble the disulfide of interest, we used a bottom-up method to acquire fragmentation intensity data for the peptide of interest (Figure S5). MS/MS analysis of the MMADHC peptide spanning residues 251-263 was extracted for underivatized, carboxyamidomethyl (CAM)-modified or N-ethylmaleimide (NEM)-modified or both CAM and NEM modifications in the tryptic peptide. The sums of the total ion count for all the detected peaks for each derivatized form was computed and tabulated (Table S1). No underivatized cysteines were detected. In the sample containing ^ΔN^MMADHC-S-cob(III) and ^ΔN^MTR, the CAM-CAM modification comprised ∼90% of the total ion count, consistent with the presence of a disulfide bond. The control sample, lacking ^ΔN^MTR, contained a mixture of CAM-NEM and NEM-CAM modifications (∼33% each), which is consistent with protection of Cys-261 or Cys-262 by B_12_ in the ^ΔN^MMADHC-S-cob(III) complex (Figure S5). The remaining ∼30% of the sample was 2xCAM modified, revealing the presence of a disulfide-containing peptide, which likely formed during sample preparation due to weak cobalamin coordination and the ability of the vicinal cysteine to displace it.

As an additional control, we tested the C261S/C262S double mutant in the full-length MMADHC background, containing the N-terminal intrinsically disordered segment. Unexpectedly, H_2_OCbl bound to C261S/C262S MMADHC with a *K*_D_ (8.4 ± 1.9 µM) that is comparable to that seen with wild-type MMADHC, suggesting an alternate B_12_-binding site. However, cofactor loaded C261S/C262S MMADHC was unable to transfer to ^ΔN^MTR (Figure S6A), unlike the wild-type full-length protein (Figure S6B). The C262S mutant in the full-length MMADHC background formed a complex with ^ΔN^MTR (Figure S6C), suggesting that the C-terminal B_12_ site is functional even when the alternate site N-terminal one is loaded with cofactor.

The relevance of the cofactor loading mechanism was tested with clinical variants of MMADHC variants (T182N, L259P, Y249C and D246G) (Figure 1E), correlated with homocystinuria^19, 21^. The *K*_D_ for H_2_OCbl coordinating to ^ΔN^MMADHC is 9.1 ± 1.6 µM and the *k*_obs_ is 0.060 ± 0.008 min^-1^. The variants display 2-to 4-fold lower *k*_obs_ values and *K*_D_ values compared to ^ΔN^MMADHC (Figure 5A,B). ^ΔN^MTR and ^ΔN^MMADHC form a complex in the absence of B_12_, which was observed by native PAGE and size exclusion chromatography (Figure 5C,D). While the relevance of this “pre-loading” complex is not known, the four clinical variants of ^ΔN^MMADHC destabilize it as evidenced by the variable increase in free ^ΔN^MMADHC as monitored by size exclusion chromatography (Figure 5D). The Y249C variant is most severely impacted and virtually no stable complex was observed with it while the D246G variant was minimally impacted. We speculate that this preloading ^ΔN^MTR-^ΔN^MMADHC complex might be responsible for recruiting MMACHC for cofactor loading.

**Figure 5.**
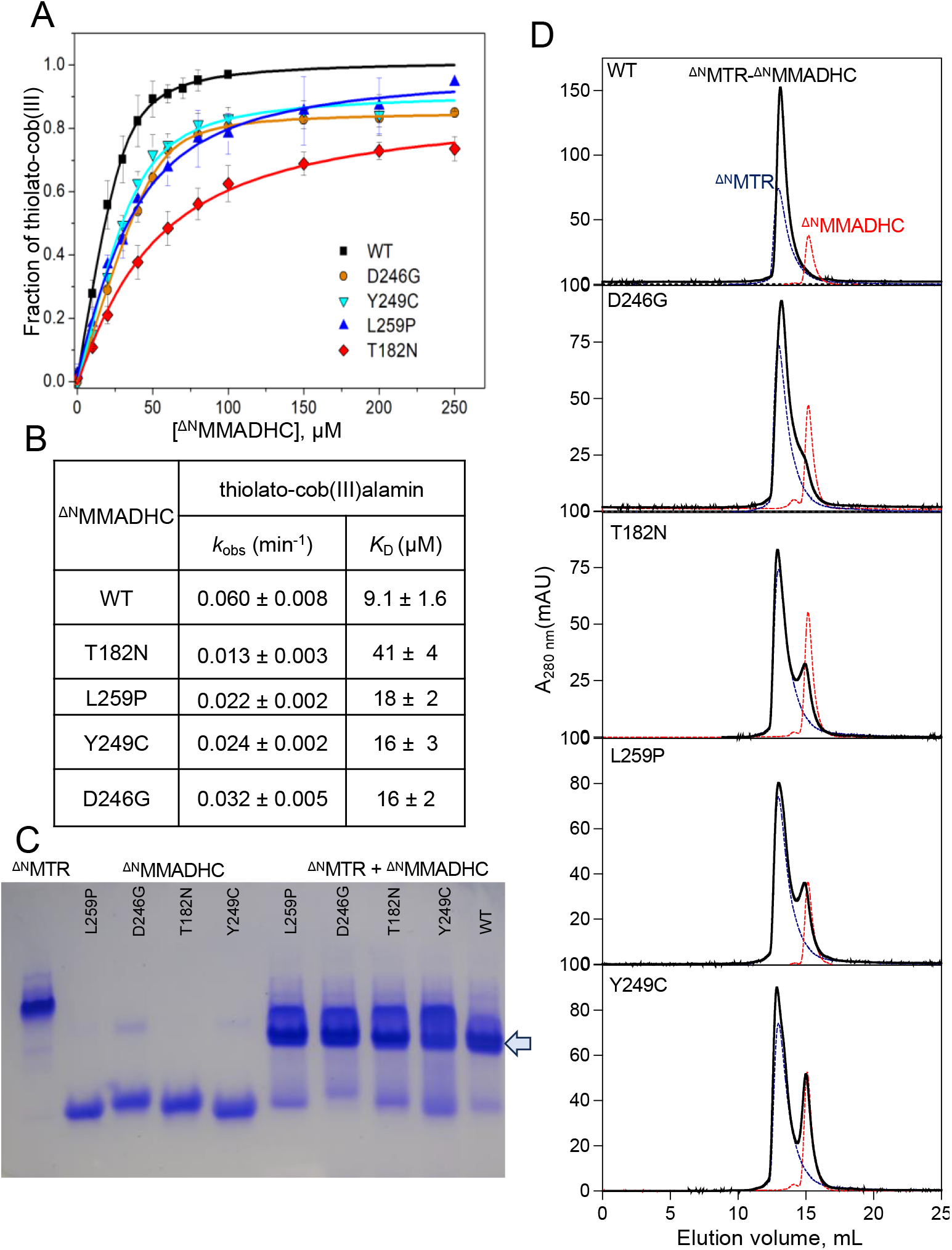
Clinical variants of MMADHC impair cobalamin binding and complex formation with MTR. **A, B.** H_2_OCbl (20 µM) was incubated with ^ΔN^MMADHC (0-100 µM) at room temperature for 1 h. The fraction of thiolato-cob(III)alamin formed was estimated from the decrease in absorbance at 351 nm (A) and the *k*_obs_ and *K*_D_ values were estimated from these traces. The data represent the mean ± SD of 3 independent experiments (B). **C**. Native PAGE analysis of the ^ΔN^MTR-^ΔN^MMADHC complexes (blue arrow). **D.** ^ΔN^MTR (20 μM) was mixed with wild-type or variant ^ΔN^MMADHC (20 μM) in Buffer A at room temperature for 10 min before separation on a size exclusion column equilibrated with the same buffer. ^ΔN^MTR (blue dashed trace) and ^ΔN^MMADHC (red dashed trace) elute at 13 and 15 mL, respectively. The ^ΔN^MTR-^ΔN^MMADHC complexes (black traces) elute between 13-13.5 mL. The data are representative of 2 independent trials.

## Discussion

Cobalt-sulfur (Co-S) coordination is rare in Nature and has been described so far in the crystal structure of the bacterial cobalamin transporter, BtuM^26^, and postulated in B_12_-dependent mercury methyltransferase HgcA^27^. MMADHC represents only the second structurally characterized example of Co-S coordination^17^, although its functional relevance has been elusive and limited understanding of the cytoplasmic branch of the B_12_ trafficking pathway. Studies on the bacterial homologs of chaperones involved in the mitochondrial branch provided early and rich insights into how the cofactor is processed and translocated in this compartment^3, 28^. In contrast, MMADHC and MMACHC are absent in bacteria, suggesting that the mechanisms for cofactor loading onto bacterial and human MTR are likely to be different. In this study, we have uncovered a central role for Co-S coordination chemistry during cofactor loading onto MTR, the cytoplasmic target of the B_12_ trafficking pathway and established its relevance with clinical variants of MMADHC, which are associated with homocystinuria.

Formation of transient protein-protein complexes between cofactor-loaded donor and apo-acceptor proteins is a common strategy in metal trafficking pathways^29^. We find that while apo-^ΔN^MMADHC and ^ΔN^MTR form a stable complex, the B_12_-bound form of ^ΔN^MMADHC engages transiently in an interprotein complex with ^ΔN^MTR, which is stabilized by mutation of Cys-262 but destabilized by each of the clinical variants of MMADHC that were tested (Figure 5C). Our data support a model for cofactor loading in which the Cys-262 thiolate functions as a nucleophile to displace and deposit cob(I)alamin on MTR, concomitantly forming a disulfide on MMADHC (Figure 4A). While the physiological relevance of the association of apo-MMADHC with MTR is not known, we speculate that it might represent a pre-loading complex needed for cofactor transfer from MMACHC to MTR. A stable MMADHC-MMACHC complex is not observed in the absence of B_12_ ^23^, highlighting differences between the affinity of apo-MMADHC for MTR versus MMACHC.

Displacement of cob(I)alamin from MMADHC leads to its concomitant transfer to MTR, which is followed by oxidation to OH_2_Cbl or conversion to MeCbl in the presence of AdoMet (Figure 4A). Since MTR has a dedicated repair system, comprising MTRR and NADPH (Figure 1A), cofactor oxidation during the loading step can be readily dealt with. In our study, a truncated form of MTR, lacking the CH_3_-H_4_F domain was used, which precluded analysis of whether AdoMet or CH_3_-H_4_F is the preferred methyl donor during cofactor loading. A series of bacterial MTR structures obtained by cryo-EM and AlphaFold2 analyses, has provided snapshots into the conformational switching between the reactivation (with cob(II)alamin bound) and resting (with MeCbl bound) states of the enzyme^10^. In the reactivation state, the AdoMet and B_12_ domains are juxtaposed, positioned for methyl group transfer (Figure 1C). However, the B_12_ domain, which is largely buried, is not readily accessible to MMADHC. A recent crystal structure of the mitochondrial B_12_ protein, MMUT, in complex with its chaperone, has captured a large 180° rotation, which exposes the B_12_ domain to solvent^30^. We speculate that a similar conformational switching of the B_12_ domain could be involved for cofactor loading onto MTR. The roles, if any, of the homocysteine and CH_3_-H_4_F modules, in regulating cofactor transfer from MMADHC is not known. CH_3_-H_4_F binding to bacterial MTR uncaps and exposes the B_12_ domain, switching the enzyme from a resting to a catalytically active conformation^10^. While it is possible CH_3_-H_4_F binding plays a similar role in increasing access to the B_12_ domain, it does not appear to be essential for cofactor loading in human MTR, since both the truncated variant lacking this domain and the full-length protein in the absence of CH_3_-H_4_F, were loaded with B_12_ by MMADHC.

Surprisingly, our study identified an alternate B_12_-binding site in full-length MMADHC, which based on the absorption spectrum, also binds B_12_ via Co-S coordination (Figure S6). There are four cysteines in the N-terminal disordered third of the protein (Cys-6, Cys-19, Cys-52 and Cys-82), which could potentially provide the sulfur ligand. None of the cysteine residues are conserved and Cys-6 and Cys-19 reside in the mitochondrial leader sequence. From the clinical mutation data, it is known that the first 116 residues in MMADHC are dispensable for MTR function^19^, and our biochemical studies reveal that B_12_ cannot be loaded from this N-terminal site onto MTR (Figure S6).

While our study describes both a role and a specific mechanism for MMADHC-dependent B_12_ transfer to human MTR, there are important gaps that remain to be filled. Co-S coordination is central to cobalamin binding to MMADHC as well as to formation of the interprotein complexes between it and MMACHC or MTR. It predicts that the Cys-261-containing face of MMADHC (Figure 1E) must interact with MMACHC and MTR in a mutually exclusive fashion. MMADHC forms a tight complex with cofactor-loaded MMACHC^17^, which is believed to be the first protein that receives and processes the cofactor in the cytoplasm^31^. It is presently unclear how cobalamin is transferred from MMACHC to MMADHC for subsequent delivery to MTR. Given the weak affinity of MMADHC for free H_2_OCbl and the low concentration of free cofactor in cells, it is unlikely that the MMADHC-S-cob(III) complex represents a free-standing intermediate in the trafficking pathway. Instead, it is more likely that the transfer from MMACHC to MTR is mediated via MMADHC in a transient complex that involves all three proteins. The chemistry underlying the escorted journey of cobalamin from its point of entry in the cytoplasm to MTR, involves intricate control of coordination chemistry and protein conformational switching. Our study fills an important piece in this puzzle by assigning a chemical function to MMADHC in cofactor loading onto MTR, and demonstrates the impact of patient mutations in corrupting the process.

## Acknowledgements

This work was supported in part by grants from the National Institutes of Health (RO1-DK45776 to R.B and K99-GM1434820 to RM). We acknowledge Dr. Uhn-Soo Cho (University of Michigan) for supporting the insect cell expression of MTR in his laboratory and thank Dr. Kazuhiro Yamada (University of Michigan) for the cDNA expressing full-length MTR. We thank the Proteomics & Metabolomics Facility (RRID:SCR_021314), Nebraska Center for Biotechnology (University of Nebraska-Lincoln) for the mass spectrometry analysis.

## Author Contributions

RM, AG and ZL performed biochemical studies. RM and MR performed EPR analysis. RM and SA cloned and purified full length MTR. JS performed MS analysis. RM and RB drafted the manuscript, and all authors edited it.

## Data and materials availability

All data are available in the manuscript or supplementary materials.

## Methods

### Materials

Cob(II)alamin was generated by photolysis of AdoCbl as previously described^32^, H_2_OCbl (Cat # 95200), MeCbl (Cat # M9756) were purchased from Sigma Aldrich. *S*-Adenosyl methionine was purchased from Cayman Chemicals, MI (Cat # 13956). Isopropyl β-;D-1-thiogalactopyranoside (IPTG) (Cat # I2481C) was from Gold Biotechnology. Ni(II)-NTA resin (Cat # 30210) was from Qiagen and amylose resin (Cat # 8021) from New England BioLabs. Primers were purchased from Integrated DNA Technologies.

### Cloning, expression and purification of ^ΔN^MTR

The gene for human MTR was codon optimized for expression in *E. coli* and cloned into a pET-28 vector between the BamH1 and Xho1 restriction sites to generate an N-terminal 6xHis tag construct with a thrombin cleavage site, pET28-thrombin-MTR. The N-terminal half of full-length MTR was truncated to span residues 661-1265, using the Quikchange kit (Qiagen) to generate pET28-thrombin-^ΔN^MTR. The sequence for the forward primer was: 5’ ATA TGG CTA TTC AGA CCG ATG AAT GGC 3’; the reverse primer was: 5’ CGG TCT GAA TAG CCA TAT GGC TGC CG 3’.

*E. coli* BL21 (DE3) cells were transformed with pET28-thrombin-^ΔN^MTR and grown overnight at 37 °C in 100 mL Luria Bertani medium (LB) containing kanamycin (50 µg•mL^-1^). 12 × 1 L of LB medium containing kanamycin (50 µg•mL^-1^) was inoculated with 10 mL of the starter culture and grown at 37 °C. After 6 h, when the OD_600_ had reached ∼0.6-0.8, the temperature was reduced to 15 °C. The cultures were induced with 50 µM IPTG, and cells were harvested 16-18 h later. Cell pellets were stored at -80 °C until use. For purification of ^ΔN^MTR, the cell pellets were suspended in 400 mL of buffer containing 50 mM Tris-HCl, pH 8.0, 500 mM KCl, 15 mM imidazole, 10 mM β-mercaptoethanol (β-ME), 0.15 mg/mL lysozyme, and 1 tablet of EDTA-free cOmplete™ Protease Inhibitor Cocktail (Roche) was added to buffer and the cell suspension was stirred at 4 °C for 40 min and sonicated (power setting = 5) on ice for 20 min at 15 sec intervals separated by 60 sec cooling periods. The sonicate was centrifuged at 38,000 × *g* for 30 min and the supernatant was loaded on a Ni-NTA agarose column (2.5 × 5 cm, Qiagen) pre-equilibrated with buffer (50 mM Tris-HCl, pH 8.0, 500 mM KCl, 15 mM imidazole). The column was washed with 200 mL wash buffer (50 mM Tris-HCl, pH 8.0, 500 mM KCl, 45 mM imidazole), and eluted with a 300 mL linear gradient, ranging from 45-400 mM imidazole in the wash buffer. The fractions containing His-^ΔN^MTR were identified by SDS-PAGE analysis, pooled, and concentrated in Amicon centrifugation devices (MWCO = 50 kDa, Sigma-Aldrich). The protein monomers were separated from oligomers by size exclusion chromatography on a Superdex 200 column (120 mL, GE Healthcare) equilibrated with buffer containing 50 mM Tris-HCl, pH 7.4, 500 mM KCl and 10% glycerol. Fractions corresponding the His_6_-^ΔN^MTR monomer were concentrated, pooled, and stored at -80 C until use.

### Cloning, expression and purification of H785A ^ΔN^MTR

The H785A mutation was introduced into the pET28-thrombin-^ΔN^MTR construct using the Quikchange kit (Qiagen). The sequence for the forward primer was: 5’ TTA AAG GTG ATG TGG CTG ATA TTG GC 3’; the reverse primer had the complementary sequence. The expression and purification of H785A His_6_-^ΔN^MTR was carried out as described above for His_6_-^ΔN^MTR.

### Cloning, expression and purification of full-length MTR

Full-length human MTR was expressed in insect cells and purified as described previously^24^. Briefly, MTR cDNA was inserted into the pFastBac Dual LIC cloning vector with an N-terminal HisX_6_-tag followed by a maltose binding protein (MBP) tag and a TEV cleavage site between MBP and MTR. Hi5 insect cells were infected with recombinant baculovirus and incubate for 44 h at 27 °C, 110 rpm. The cell pellet was suspended in buffer containing 50 mM Tris-HCl, pH 7.4, 300 mM NaCl,1 tablet of EDTA-free cOmplete Protease Inhibitor Cocktail, 0.20 mg/mL lysozyme and 3 mM β-ME. The cells were lysed by 4 cycles of freezing in liquid N_2_ and thawing in a water bath at room temperature. The cell lysates were centrifuged at 38,000 *× g* at 4°C for 2 h and the supernatant was loaded onto an amylose resin column (2.5 × 5 cm) equilibrated with buffer (50 mM Tris-HCl pH 8, 300 mM NaCl and 10% glycerol) at 4 °C. His-MBP-MTR was eluted with 50 mM Tris-HCl pH 8, 300 mM NaCl, 10% glycerol and 10 mM maltose. Fractions containing His-MBP-MTR were pooled and concentrated using an Amicon Ultra-15 (MWCO=50 kDa) filter. Protein concentration was estimated by the Bradford method using the Bio-Rad protein assay reagent, flash frozen in liquid N_2_ and stored at -80°C until use.

### Expression and purification of ^ΔN^MMADHC variants

MMADHC lacking the first 115 residues (^ΔN^MMADHC) was purified as described previously using the pGEX-MMADHC construct^17^. Mutations were introduced using the Quick-change kit (Qiagen). The sequences of the forward primers are noted below; the reverse primers had complementary sequences.

T182N: 5′-ACTGTAACACAAAAAAATAAGATGATATGACT-3′, L259P: 5′-TTCTCTGTTGATGACCCGGGATGCTGTAAAGTG-3′ Y249C: 5′-GAAACTGATAACGCTGCCGACATTTAGGATTC-3′ D246G: 5′-ACTCTTTTTGAAACTGGTGAACGCTACCGACAT-3′ C261/C262S: 5’-TCCTCTAAAGTGATTCGTCATAGTCTCTGG-3’

### Expression and purification of MMACHC

A previously reported truncated form of MMACHC (1-244) with a C-terminal His_6_-tag which was expressed and purified for assays as described previously^33^.

### Buffer conditions for experiments

Unless otherwise noted, all titrations and kinetic experiments were performed in Buffer A containing 0.1 M HEPES, pH 7.4, 150 mM KCl, and 10% glycerol.

### Formation of ^ΔN^MMADHC-S-cob(III)alamin

The ^ΔN^MMADHC-S-cob(III) complex was prepared as described previously^17^ by mixing OH_2_Cbl (100 µM) with a slight excess of ^ΔN^MMADHC (120 µM) in Buffer A at room temperature for 1 h. Unreacted H_2_OCbl was removed by centrifugation using a NanoSep filter device (MWCO=10 kDa, PALL). The C262S ^ΔN^MMADHC-S-cob(III) and C261S ^ΔN^MMADHC-S-cob(III) complexes were prepared similarly.

### Cofactor transfer from ^ΔN^MMADHC to ^ΔN^MTR

The ^ΔN^MMADHC-S-cob(III) complex (20 µM) was incubated with ^ΔN^MTR (40 µM) in the Buffer A at 20°C. The spectra were monitored every minute for 50 min until no further changes were observed. The absorbance changes at 350 nm or 555 nm were fitted to a double exponential decay using GraphPad Prism Software. The reaction mixture (150 µL) was then centrifuged at 21,000 ×*g* at 4 °C for 10 min and 100 µL of the supernatant was loaded on to a Superdex 200 column (25 mL, GE Healthcare) equilibrated with the Buffer A, at 4 °C at a flow rate of 0.2 mL/min. The elution profile was recorded at 280 and 350 nm. Under these conditions, the elution volumes for ^ΔN^MTR and ^ΔN^MMADHC were, 13 and 16 mL, respectively. Cofactor transfer from C261S or C262S ^ΔN^MMADHC-S-cob(III) complexes to ^ΔN^MTR were tested similarly.

### Cofactor transfer from ^ΔN^MMADHC to ^ΔN^MTR under anaerobic conditions

The ^ΔN^MMADHC-S-cob(III) complex was prepared as described above and exchanged into anaerobic Buffer A in the anaerobic chamber (< 0.5 ppm O_2_). A 100 μL stock solution of ^ΔN^MTR was purged with N_2_ gas on ice for 45 min in a sealed 1.5 mL Eppendorf tube before moving it into the anaerobic chamber. To monitor cofactor transfer, ^ΔN^MMADHC-S-cob(III) (20 µM) was mixed with ^ΔN^MTR (40 µM) in Buffer A at room temperature under anaerobic conditions. Spectra were monitored every 30 s for 50 min until no further changes were observed. The same experiment was repeated in the presence of 500 μΜ ΑdoMet. The maximal absorbance change was recorded at 570 nm and fitted to a double exponential decay using GraphPad Prism Software. The concentration of MeCbl transferred was determined using Δε_570 nm_ of 3.5 mM^-1^ cm^-1^, which was calculated from ΔA (^ΔN^MMADHC-S-cob(III) – ^ΔN^MTR•MeCbl). To confirm cofactor transfer to MTR, the reaction mixture was analyzed by analytical size exclusion chromatography on a Superdex 200 column (25 mL, GE Healthcare) as described above.

### Analysis of C262S ^ΔN^MMADHC-S-cob(III)-^ΔN^MTR complex

C262S ^ΔN^MMADHC-S-cob(III) (25 µM) was incubated with varying concentrations of ^ΔN^MTR (0-60 µM) in Buffer A at 20°C. Spectra were recorded every 30 min until no further changes were observed. The absorbance change at 555 nm was fitted to the equation:

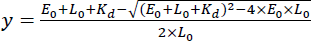

where E_0_ and L_0_ are the total concentration of ^ΔN^MTR and C262S ^ΔN^MMADHC-S-cob(III), respectively.

### Interaction of ^ΔN^MMADHC clinical variants with H_2_OCbl-^ΔN^

MMADHC variants (40 µM) were mixed with H_2_OCbl (20 µM) in Buffer A at room temperature (23 °C). The UV-visible spectrum was recorded every minute for 1 h to monitor reaction progress. The change in absorbance at 351 nm (Δe_251nm_=11.8 mM^-1^cm^-1, 33^) was fitted to a single exponential equation:

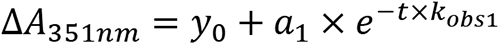

To estimate the affinity of ^ΔN^MMADHC for H_2_OCbl, H_2_OCbl (20 µM) was incubated with 0-100 µM ^ΔN^MMADHC for 1 h at room temperature. The resulting concentrations of ^ΔN^MMADHC-thiolato-cob(III) was calculated from the A_351nm_ and the fraction of ^ΔN^MMADHC-S-cob(III) (y) was fitted to the equation:

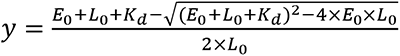

where E_0_ and L_0_ are the total concentration of ^ΔN^MMADHC and H_2_OCbl, respectively.

### Complex formation between clinical variants of apo-^ΔN^MMADHC and apo-^ΔN^MTR

^ΔN^MMADHC variants (50 µM) were mixed with ^ΔN^MTR (50 µM) in Buffer A (15 μL) at room temperature for 5 min. Sample were mixed with 15 μL 2X native gel loading buffer (62.5 mM Tris-HCl, pH 6.8, 40% glycerol, 0.01% bromophenol blue) and 10 μL samples were loaded onto a 4-15 % gradient gel (Bio-Rad) and separated using running buffer (2.5mM Tris, 19.2 mM glycine, pH 8.3) at 4 °C, 100 V for 160 min. For gel filtration analyses, 20 μM ^ΔN^MTR was mixed with 20 μM wild-type or variant ^ΔN^MMADHC in Buffer A at room temperature for 10 min before separation on a BioRad enrich-650 size exclusion column equilibrated with the same buffer. ^ΔN^MTR eluted at 13 mL, ^ΔN^MMADHC eluted at 15 mL. The ^ΔN^MTR-^ΔN^MMADHC complexes eluted between 13-13.5 mL.

### HPLC analysis of cobalamins

For the reactivation reaction, 40 µM ^ΔN^MTR, 1 mM AdoMet, 20 µM cob(II)alamin, 300 µM NADPH and 40 µM MTRR were mixed in Buffer A under anaerobic conditions for 60 min. The reaction was quenched with 2% trifluoroacetic acid and centrifuged for 10 min at 13,600 x *g*. For the transfer reaction, ^ΔN^MMADHC-S-cob(III) complex (20 µM) was mixed with ^ΔN^MTR (40 µM) in Buffer A containing 500 μM AdoMet under anaerobic conditions. UV-visible spectra were monitored for 30 min every 30 sec until no additional changes were observed. The reaction was quenched with 2% trifluoroacetic acid, and centrifuged for 10 min at 13,600 x *g.* The supernatant (100 µl) was injected onto an Alltima HP C18 column (Grace, 250mm x 4.6 mm, 5 µM) at a flow rate of 1 ml min^-1^. The mobile phase consisted of solvent A: 0.1 % trifluoroacetic acid in water and solvent B: 0.1 % trifluoroacetic acid in acetonitrile. Samples were eluted at a flow rate of 1 ml min^-1^ with a gradient from 8-32% solvent B for 35 min, and the absorbance was monitored at 254 nm and 350 nm. The standards MeCbl (20 µM) and OH_2_Cbl (20 µM) were treated similarly.

### EPR spectroscopy

EPR spectra were recorded on a Bruker EMX 300 equipped with a Bruker 4201 cavity and a ColdEdge cryostat. The temperature was controlled by an Oxford Instruments MercuryiTC temperature controller. EPR spectra were recorded at 80 K using the following parameters: 9.37 GHz microwave frequency, power 2 mW, modulation amplitude 10 G, modulation frequency 100 kHz, 3000 G sweep width centered at 3500 G, conversion time 164 ms, time constant 82 ms. Five scans were collected per measurement. EPR simulations were performed using the EasySpin program^34^. The ^ΔN^MTR-cob(II)alamin spectrum was simulated (Figure S2E) as a mixture of 45% His-coordinated cob(II)alamin and 55% H2O-coordinated cob(II)alamin, using the following parameters: His-coordinated: *g*x=2.296, *g*y=2.225, *g*z=1.999; principal hyperfine coupling constants, Ax=17, Ay=16, Az=303 MHz; AN = 48 MHz; H2O-coordinated: principal *g*-values, *g*x=2.402, *g*y=2.309, *g*z=1.991; principal hyperfine coupling constants, Ax=228, Ay=203, Az=384 MHz. The ^ΔN^MTR^H785A^-cob(II)alamin spectrum was simulated (Figure S2F), using the following parameters: principal *g*-values, *g*x=2.397, *g*y=2.304, *g*z=1.994; principal hyperfine coupling constants, Ax=223, Ay=192, Az=382 MHz.

Samples were prepared in an anaerobic chamber as follows. ^ΔN^MTR (150 μΜ) or ^ΔN^MTR H785A (150 μΜ) were mixed with cob(II)alamin (100 µM) in Buffer A, in a total volume of 250 µL. The samples were incubated for 15 min at room temperature, transferred to EPR tubes and frozen in liquid nitrogen. The base-on cob(II)alamin control sample was prepared in the same buffer, and the base-off sample was adjusted to pH 2.0 with concentrated HCl.

### Mass spectrometry

The strategy for MS sample preparation is outlined in Figure S5. Briefly, ^ΔN^MMADHC-S-cob(III) (20 μΜ) was prepared as described above, mixed with 30 μΜ ^ΔN^ΜΤR in Buffer A and incubated for ∼ 1 h at room temperature in a reaction volume of 250 μL. Then, 0.6 mM N-ethylmaleimide (NEM) was added and the reaction mixture, protected from light, was incubated for 2 h at room temperature. Unreacted NEM was removed by centrifugation using a NanoSep filter device (MWCO=10 kDa, PALL) and washed 6 times in buffer containing 100 mM ammonium bicarbonate, pH 8. The sample was concentrated to ∼100 μL and treated with 5 μL 2% SDS before the second alkylation step. The sample was reduced with 10 mM Tris(2-carboxyethyl) phosphine hydrochloride (TCEP•HCl) and incubated at 55 °C for 1 h. Then, 5 μL 375 mM iodoacetamide was added to the samples and incubated for 30 min protected from light. The samples were flash frozen in liquid N_2_ until further use. The control ^ΔN^MMADHC-S-cob(III) complex (20 μΜ) was treated in an identical fashion in parallel.

The proteins were separated on a denaturing SDS/PAGE gel. The band at ∼20 kDa corresponding to ^ΔN^MMADHC was excised and washed with 0.1 M ammonium bicarbonate in 10% acetonitrile to remove SDS and the Coomassie blue stain. The samples were digested at 37 °C overnight with trypsin (15 µg/mL w/v) and extracted with 3 washes with 50% acetonitrile in 50 mM ammonium bicarbonate. The extracted peptides were combined, concentrated by evaporation, dissolved in 50 μL 0.1% formic acid and analyzed after a 1 h run on a 0.075 mm x 250 mm C18 Waters CSH column interfaced with an Orbitrap Eclipse mass spectrometer (Thermo Scientific, Waltham, Massachusetts, US). The instrument operated in data-dependent acquisition mode to perform bottom-up proteomics and the generated data were analyzed with Mascot (Matrix Science, London, UK; version 2.7.0). The program was set up to search the cRAP_20150130.fasta (125 common contaminant sequences), uniprot-human_20210508 (77,027 entries) databases for tryptic peptides. Mascot was searched with a fragment ion mass tolerance of 0.060 Da and a parent ion tolerance of 10.0 ppm. Deamidation of asparagine and glutamine, oxidation of methionine, carbamidomethylation (CAM), NEM and NEM + water derivatives of cysteine and lysine were specified in Mascot as variable modifications.

Scaffold (version Scaffold_4.8.9, Proteome Software Inc., Portland, OR) was used to validate the MS/MS based peptide (and protein) identification. Peptide identification was accepted if it met a >80.0% probability threshold using the Peptide Prophet algorithm ^45^ with Scaffold delta-mass correction. The peptide lists for MMADHC were exported into Excel, which was used to sort out the frequency of peptide modifications for the tryptic peptide spanning residues 251-263. The sum of total ion count as computed for each modification and the percentages were calculated based on the sum of the total ion count.

## References

(1) Banerjee, R.; Gouda, H.; Pillay, S. Redox-Linked Coordination Chemistry Directs Vitamin B_12_ Trafficking. Acc Chem Res 2021, 54 (8), 2003–2013. DOI: 10.1021/acs.accounts.1c00083. Gherasim, C.; Lofgren, M.; Banerjee, R. Navigating the B_12_ road: assimilation, delivery and disorders of cobalamin. J Biol Chem 2013, 288, 13186–13193. DOI: 10.1074/jbc.R113.458810.

(2) Watkins, D.; Rosenblatt, D. S. Inborn errors of cobalamin absorption and metabolism. Am J Med Genet C Semin Med Genet 2011, *157C* (1), 33–44. DOI: 10.1002/ajmg.c.30288.

(3) Padovani, D.; Labunska, T.; Palfey, B. A.; Ballou, D. P.; Banerjee, R. Adenosyltransferase tailors and delivers coenzyme B_12_. Nat Chem Biol 2008, 4 (3), 194–196. DOI: nchembio.67 [pii] 10.1038/nchembio.67.

(4) Ruetz, M.; Campanello, G. C.; McDevitt, L.; Yokom, A. L.; Yadav, P. K.; Watkins, D.; Rosenblatt, D. S.; Ohi, M. D.; Southworth, D. R.; Banerjee, R. Allosteric Regulation of Oligomerization by a B_12_ Trafficking G-Protein Is Corrupted in Methylmalonic Aciduria. Cell Chem Biol 2019, 26 (7), 960–969 e964. DOI: 10.1016/j.chembiol.2019.03.014. Campanello, G. C.; Ruetz, M.; Dodge, G. J.; Gouda, H.; Gupta, A.; Twahir, U. T.; Killian, M. M.; Watkins, D.; Rosenblatt, D. S.; Brunold, T. C.;, et al. Sacrificial Cobalt-Carbon Bond Homolysis in Coenzyme B_12_ as a Cofactor Conservation Strategy. J Am Chem Soc 2018, 140 (41), 13205–13208. DOI: 10.1021/jacs.8b08659. Gouda, H.; Mascarenhas, R.; Pillay, S.; Ruetz, M.; Koutmos, M.; Banerjee, R. Patient mutations in human ATP:cob(I)alamin adenosyltransferase differentially affect its catalytic versus chaperone functions. J Biol Chem 2021, 297 (6), 101373. DOI: 10.1016/j.jbc.2021.101373. Ruetz, M.; Campanello, G. C.; Purchal, M.; Shen, H.; McDevitt, L.; Gouda, H.; Wakabayashi, S.; Zhu, J.; Rubin, E. J.; Warncke, K.;, et al. Itaconyl-CoA forms a stable biradical in methylmalonyl-CoA mutase and derails its activity and repair. Science 2019, 366 (6465), 589–593. DOI: 10.1126/science.aay0934. Gouda, H.; Mascarenhas, R.; Ruetz, M.; Yaw, M.; Banerjee, R. Bivalent molecular mimicry by ADP protects metal redox state and promotes coenzyme B(12) repair. Proc Natl Acad Sci U S A 2023, 120 (11), e2220677120. DOI: 10.1073/pnas.2220677120 From NLM Medline.

(5) Banerjee, R. V.; Matthews, R. G. Cobalamin-dependent methionine synthase. FASEB J. 1990, 4, 1450–1459.

(6) Banerjee, R.; Frasca, V.; Ballou, D. P.; Matthews, R. G. Participation of cob(I)alamin in the reaction catalyzed by methionine synthase from *Escherichia coli:* A steady-state and rapid reaction kinetic analysis. Biochemistry 1990, 29, 11101–11109. Drummond, J. T.; Huang, S.; Blumenthal, R. M.; Matthews, R. G. Assignment of enzymatic function to specific protein regions of cobalamin-dependent methionine synthase from Escherichia coli. Biochemistry 1993, 32 (36), 9290–9295.

(7) Banerjee, R. V.; Harder, S., R.; Ragsdale, S. W.; Matthews, R. G. Mechanism of reductive activation of cobalamin-dependent methionine synthase: an electron paramagnetic resonance spectroelectrochemical study. Biochemistry 1990, 29, 1129–1137.

(8) Yamada, K.; Gravel, R. A.; Toraya, T.; Matthews, R. G. Human methionine synthase reductase is a molecular chaperone for human methionine synthase. Proc Natl Acad Sci U S A 2006, 103 (25), 9476–9481.

(9) Gulati, S.; Chen, Z.; Brody, L. C.; Rosenblatt, D. S.; Banerjee, R. Defects in auxiliary redox proteins lead to functional methionine synthase deficiency. J Biol Chem 1997, 272 (31), 19171–19175.

(10) Watkins, M. B.; Wang, H.; Burnim, A.; Ando, N. Conformational switching and flexibility in cobalamin-dependent methionine synthase studied by small-angle X-ray scattering and cryoelectron microscopy. Proc Natl Acad Sci U S A 2023, 120 (26), e2302531120. DOI: 10.1073/pnas.2302531120 From NLM Medline.

(11) Datta, S.; Koutmos, M.; Pattridge, K. A.; Ludwig, M. L.; Matthews, R. G. A disulfide-stabilized conformer of methionine synthase reveals an unexpected role for the histidine ligand of the cobalamin cofactor. Proc Natl Acad Sci U S A 2008, 105 (11), 4115–4120. DOI: 10.1073/pnas.0800329105. Bandarian, V.; Pattridge, K. A.; Lennon, B. W.; Huddler, D. P.; Matthews, R. G.; Ludwig, M. L. Domain alternation switches B(12)-dependent methionine synthase to the activation conformation. Nat Struct Biol 2002, 9 (1), 53–56. DOI: 10.1038/nsb738.

(12) Lerner-Ellis, J. P.; Tirone, J. C.; Pawelek, P. D.; Dore, C.; Atkinson, J. L.; Watkins, D.; Morel, C. F.; Fujiwara, T. M.; Moras, E.; Hosack, A. R.;, et al. Identification of the gene responsible for methylmalonic aciduria and homocystinuria, *cbl*C type. Nat. Genet. 2006, 38 (1), 93–100.

(13) Kim, J.; Hannibal, L.; Gherasim, C.; Jacobsen, D. W.; Banerjee, R. A human vitamin B_12_ trafficking protein uses glutathione transferase activity for processing alkylcobalamins. J Biol Chem 2009, 284 (48), 33418–33424. DOI: M109.057877 [pii]10.1074/jbc.M109.057877.

(14) Mascarenhas, R.; Li, Z.; Gherasim, C.; Ruetz, M.; Banerjee, R. The human B12 trafficking protein CblC processes nitrocobalamin. J Biol Chem 2020, 295 (28), 9630–9640. DOI: 10.1074/jbc.RA120.014094.

(15) Kim, J.; Gherasim, C.; Banerjee, R. Decyanation of vitamin B_12_ by a trafficking chaperone. Proc. Natl. Acad. Sci. U.S.A. 2008, 105 (38), 14551–14554.

(16) Li, Z.; Gherasim, C.; Lesniak, N. A.; Banerjee, R. Glutathione-dependent One-electron Transfer Reactions Catalyzed by a B12 Trafficking Protein. J Biol Chem 2014, 289 (23), 16487–16497. DOI: 10.1074/jbc.M114.567339.

(17) Li, Z.; Mascarenhas, R.; Twahir, U. T.; Kallon, A.; Deb, A.; Yaw, M.; Penner-Hahn, J.; Koutmos, M.; Warncke, K.; Banerjee, R. An Interprotein Co-S Coordination Complex in the B12-Trafficking Pathway. J Am Chem Soc 2020, 142 (38), 16334–16345. DOI: 10.1021/jacs.0c06590.

(18) Yamada, K.; Gherasim, C.; Banerjee, R.; Koutmos, M. Structure of Human B_12_ Trafficking Protein CblD Reveals Molecular Mimicry and Identifies a New Subfamily of Nitro-FMN Reductases. J Biol Chem 2015, 290 (49), 29155–29166. DOI: 10.1074/jbc.M115.682435. Froese, D. S.; Kopec, J.; Fitzpatrick, F.; Schuller, M.; McCorvie, T. J.; Chalk, R.; Plessl, T.; Fettelschoss, V.; Fowler, B.; Baumgartner, M. R.;, et al. Structural Insights into the MMACHC-MMADHC Protein Complex Involved in Vitamin B12 Trafficking. J Biol Chem 2015, 290 (49), 29167–29177. DOI: 10.1074/jbc.M115.683268.

(19) Coelho, D.; Suormala, T.; Stucki, M.; Lerner-Ellis, J. P.; Rosenblatt, D. S.; Newbold, R. F.; Baumgartner, M. R.; Fowler, B. Gene identification for the cblD defect of vitamin B_12_ metabolism. N Engl J Med 2008, 358 (14), 1454–1464. DOI: 358/14/1454 [pii] 10.1056/NEJMoa072200.

(20) Stucki, M.; Coelho, D.; Suormala, T.; Burda, P.; Fowler, B.; Baumgartner, M. R. Molecular mechanisms leading to three different phenotypes in the cblD defect of intracellular cobalamin metabolism. Hum Mol Genet 2012, 21 (6), 1410–1418, Research Support, Non-U.S. Gov’t. DOI: 10.1093/hmg/ddr579.

(21) Suormala, T.; Baumgartner, M. R.; Coelho, D.; Zavadakova, P.; Koich, V.; Koch, H. G.; Berghauser, M.; Wraith, J. E.; Burlina, A.; Sewell, A.;, et al. The *cbl*D defect causes either isolated or combined deficiency of methylcobalamin and adenosylcobalamin synthesis. J Biol Chem 2004, (41), 42742–42749.

(22) Deme, J. C.; Miousse, I. R.; Plesa, M.; Kim, J. C.; Hancock, M. A.; Mah, W.; Rosenblatt, D. S.; Coulton, J. W. Structural features of recombinant MMADHC isoforms and their interactions with MMACHC, proteins of mammalian vitamin B_12_ metabolism. Mol Genet Metab 2012, 107 (3), 352–362. DOI: 10.1016/j.ymgme.2012.07.001.

(23) Gherasim, C.; Hannibal, L.; Rajagopalan, D.; Jacobsen, D. W.; Banerjee, R. The C-terminal domain of CblD interacts with CblC and influences intracellular cobalamin partitioning Biochemie 2013, 95, 1023–1032.

(24) Mascarenhas, R.; Gouda, H.; Ruetz, M.; Banerjee, R. Human B_12_-dependent enzymes: Methionine synthase and Methylmalonyl-CoA mutase. Methods Enzymol 2022, 668, 309–326. DOI: 10.1016/bs.mie.2021.12.012 From NLM Medline.

(25) Li, Z.; Shanmuganathan, A.; Ruetz, M.; Yamada, K.; Lesniak, N. A.; Krautler, B.; Brunold, T. C.; Koutmos, M.; Banerjee, R. Coordination chemistry controls the thiol oxidase activity of the B_12_-trafficking protein CblC. J Biol Chem 2017, 292 (23), 9733–9744. DOI: 10.1074/jbc.M117.788554.

(26) Rempel, S.; Colucci, E.; de Gier, J. W.; Guskov, A.; Slotboom, D. J. Cysteine-mediated decyanation of vitamin B_12_ by the predicted membrane transporter BtuM. Nat Commun 2018, 9 (1), 3038. DOI: 10.1038/s41467-018-05441-9.

(27) Zhou, J.; Riccardi, D.; Beste, A.; Smith, J. C.; Parks, J. M. Mercury methylation by HgcA: theory supports carbanion transfer to Hg(II). Inorg Chem 2014, 53 (2), 772–777. DOI: 10.1021/ic401992y From NLM Medline.

(28) Padovani, D.; Banerjee, R. Assembly and protection of the radical enzyme, methylmalonyl-CoA mutase, by its chaperone. Biochemistry 2006, 45 (30), 9300–9306. Padovani, D.; Banerjee, R. A G-protein editor gates coenzyme B_12_ loading and is corrupted in methylmalonic aciduria. Proc Natl Acad Sci U S A 2009, 106 (51), 21567–21572. DOI: 0908106106[pii]10.1073/pnas.0908106106. Padovani, D.; Banerjee, R. A Rotary Mechanism for Coenzyme B_12_ Synthesis by Adenosyltransferase. Biochemistry 2009, 48, 5350–5357. Padovani, D.; Labunska, T.; Banerjee, R. Energetics of interaction between the G-protein chaperone, MeaB and B12-dependent methylmalonyl-CoA mutase. J. Biol. Chem. 2006, 281, 17838–17844. Mascarenhas, R.; Ruetz, M.; McDevitt, L.; Koutmos, M.; Banerjee, R. Mobile loop dynamics in adenosyltransferase control binding and reactivity of coenzyme B_12_. Proc Natl Acad Sci U S A 2020, 117 (48), 30412–30422. DOI: 10.1073/pnas.2007332117. Lofgren, M.; Banerjee, R. Loss of allostery and coenzyme B_12_ delivery by a pathogenic mutation in adenosyltransferase. Biochemistry 2011, 50 (25), 5790–5798. DOI: 10.1021/bi2006306. Lofgren, M.; Koutmos, M.; Banerjee, R. Autoinhibition and signaling by the switch II motif in the G-protein chaperone of a radical B_12_ enzyme. J Biol Chem 2013, 288 (43), 30980–30989. DOI: 10.1074/jbc.M113.499970. Lofgren, M.; Padovani, D.; Koutmos, M.; Banerjee, R. A switch III motif relays signaling between a B_12_ enzyme and its G-protein chaperone. Nat Chem Biol 2013, 9 (9), 535–539. DOI: 10.1038/nchembio.1298.

(29) Camponeschi, F.; Banci, L. Metal cofactors trafficking and assembly in the cell: a molecular view. Pure App Chem 2019, 91, 231–245.

(30) Mascarenhas, R.; Ruetz, M.; Gouda, H.; Heitman, N.; Yaw, M.; Banerjee, R. Architecture of the human G-protein-methylmalonyl-CoA mutase nanoassembly for B(12) delivery and repair. Nat Commun 2023, 14 (1), 4332. DOI: 10.1038/s41467-023-40077-4 From NLM Medline.

(31) Banerjee, R. B_12_ trafficking in mammals: A case for coenzyme escort service. ACS Chem Biol 2006, 1 (3), 149–159. DOI: 10.1021/cb6001174.

(32) Li, Z.; Kitanishi, K.; Twahir, U. T.; Cracan, V.; Chapman, D.; Warncke, K.; Banerjee, R. Cofactor Editing by the G-protein Metallochaperone Domain Regulates the Radical B_12_ Enzyme IcmF. J Biol Chem 2017, 292 (10), 3977–3987. DOI: 10.1074/jbc.M117.775957.

(33) Koutmos, M.; Gherasim, C.; Smith, J. L.; Banerjee, R. Structural basis of multifunctionality in a vitamin B_12_-processing enzyme. J Biol Chem 2011, 286 (34), 29780–29787, Research Support, N.I.H., Extramural. DOI: 10.1074/jbc.M111.261370.

(34) Stoll, S.; Schweiger, A. EasySpin, a comprehensive software package for spectral simulation and analysis in EPR. J Magn Reson 2006, 178 (1), 42–55. DOI: 10.1016/j.jmr.2005.08.013.

