## Supplementary Information for "Cobalt-sulfur coordination chemistry drives B_12_ loading onto methionine synthase"

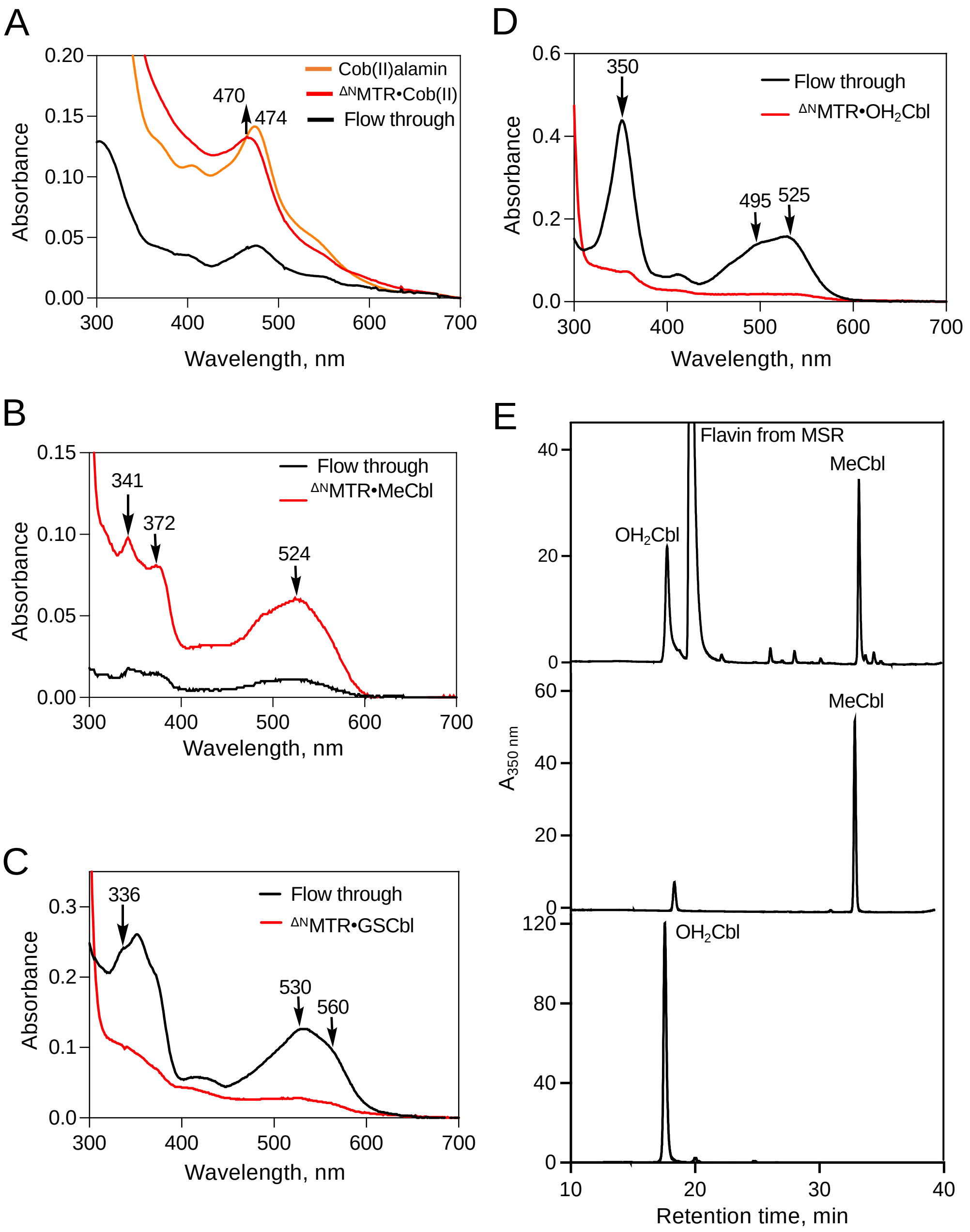


**Figure S1. MTR binds select cobalamin derivatives from solution. A.** Cob(II)alamin (15 μM, *orange*) was mixed with 30 μM $^{\Delta N}$MTR in Buffer A at 20°C under anaerobic conditions and the spectrum were recorded after 30 min (red). Most of the cobalamin remained bound to MTR following separation on a Nanosep column. The blue shift in the spectrum from 474 nm to 470 nm is consistent with cob(II)alamin binding to $^{\Delta N}$MTR. **B, C, D.** $^{\Delta N}$MTR (40 μM) was mixed with MeCbl, GSCbl or $OH_2$Cbl (20 μM each) in Buffer A at room temperature for 15 min. The samples were diluted 2X with Buffer A and centrifuged in a Nanosep column to separate unbound cobalamin. Comparison of the spectra of the flow-through (*black*) and $^{\Delta N}$MTR *(red*) fractions reveals that MeCbl binds to $^{\Delta N}$MTR (B) while GSCbl (C) and $OH_2$Cbl (D) do not. **E.** An anaerobic solution containing $^{\Delta N}$MTR (40 μM), 1 mM AdoMet, 20 μM cob(II)alamin, 300 μM NADPH and 40 μM MTRR in Buffer A was incubated for 60 min after which the reaction was quenched with 2% TFA. MeCbl formation was confirmed by HPLC analysis of the reaction mixture.

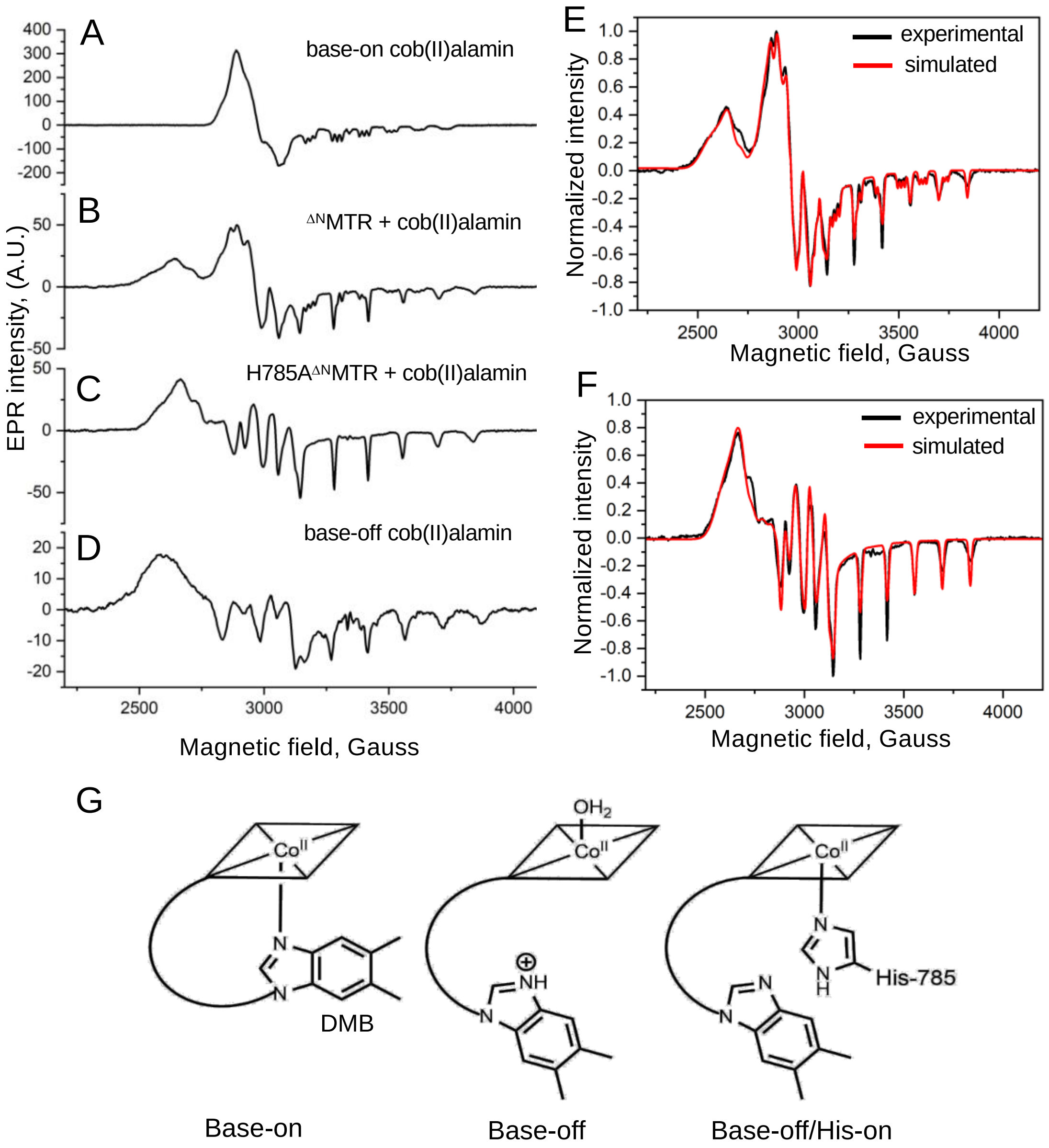


**Figure S2. EPR spectrum of $^{\Delta N}$MTR bound cob(II)alamin. A.** EPR spectra of cob(II)alamin (100 µM) in Buffer A. **B**. When $^{\Delta N}$MTR (150 µM) was added to A, a mixture of cob(II)alamin with a nitrogen (His) or oxygen ($H_2O$) ligand was obtained. **C**. Addition of $^{\Delta N}$MTR$^{H785A}$ (150 µM) to A resulted in cob(II)alamin species with an oxygen ligand. **D.** The spectrum of base-off cobalamin (100 µM) in Buffer A acidified with concentrated HCl. EPR spectra were recorded at 80 K using the following parameters: 9.37 GHz microwave frequency, 20 mW power, 10 G modulation amplitude, 100 kHz modulation frequency, 3000 G sweep width centered at 3500 G, conversion time 164 msec, time constant 82 msec. Five scans were collected for each sample. A.U. denotes arbitrary units. **E.** The experimental spectrum of $^{\Delta N}$MTR-cob(II)alamin (black) is overlaid on the simulated spectrum (red) generated as a mixture of 45% His-on cob(II)alamin and 55% base-off ($H_2O$-coordinated) cob(II)alamin. **F.** The experimental spectrum (black) of $^{\Delta N}$MTR$^{H785A}$-cob(II)alamin is overlaid on the simulated spectrum (red). The parameters used to simulate spectra in E and F are described under Methods **G**. Cartoons of 5-coordinate cob(II)alamin in the base-on, water-coordinated base-off and His-on/base-off states.

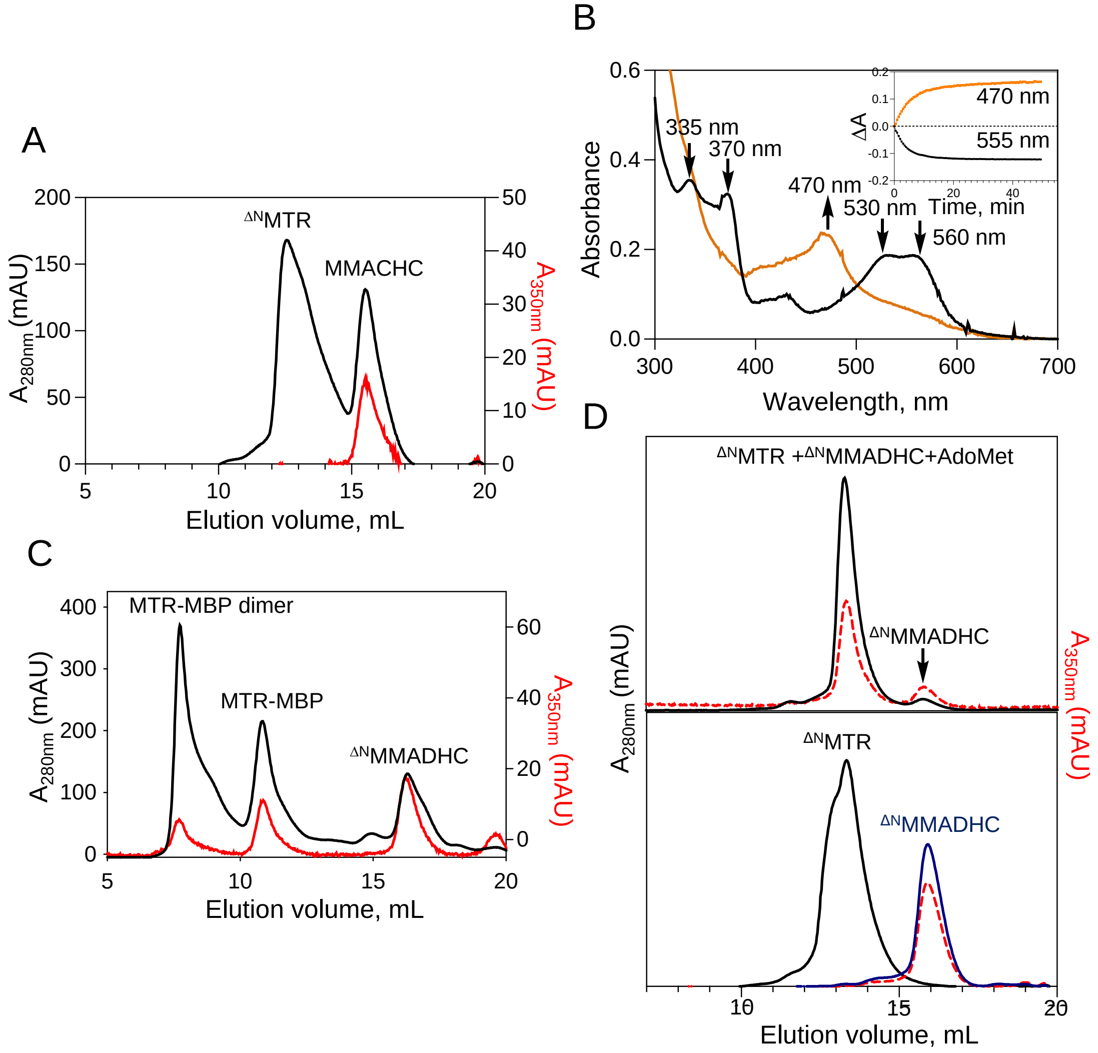


**Figure S3. $B_{12}$ transfer to MTR**. Elution profile of sample in which anaerobic cob(II)alamin (20 μM) was mixed with MMACHC (30 μM) under anaerobic conditions, and then mixed with $^{\Delta N}$MTR (40 μM) and loaded onto an analytical S200 column equilibrated with Buffer A. Protein (280 nm, black) and cobalamin (350 nm, red) were monitored at the indicated wavelengths. **B.** $^{\Delta N}$MMADHC-S-cob(III) (20 μM) was mixed with 40 μM $^{\Delta N}$MTR in Buffer A at 20°C under anaerobic conditions and UV/visible spectra were recorded every 30 s for 50 min. The initial (black trace) and final (orange trace) spectra are shown. The blue shift in the spectrum from the peak at 530 and 560 nm to 470 nm, is indicative of transfer to $^{\Delta N}$MTR. *Inset.* Change in absorbance at 555 and 470 nm versus time. **C.** Transfer of $B_{12}$ from $^{\Delta N}$MMADHC-S-cob(III) to full-length human MTR-MBP (maltose binding protein). $^{\Delta N}$MMADHC-S-cob(III) (20 μM, black trace) was mixed with MTR-MBP (80 μM) in Buffer A at 20°C. The sample was loaded onto an analytical S200 column equilibrated with Buffer A and the eluant was monitored at 280 nm (black trace) and 350 nm (red trace). **D.** $^{\Delta N}$MMADHC-S-cob(III)(20 μM) and 500 μM AdoMet, were mixed with $^{\Delta N}$MTR (40 μM) in Buffer A under anaerobic conditions and separated on an S200 column. The retention times for the $^{\Delta N}$MTR and $^{\Delta N}$MMADHC-S-cob(III) standards are shown. The elution profile confirms $B_{12}$ transfer to $^{\Delta N}$MTR from $^{\Delta N}$MMADHC-S-cob(III).

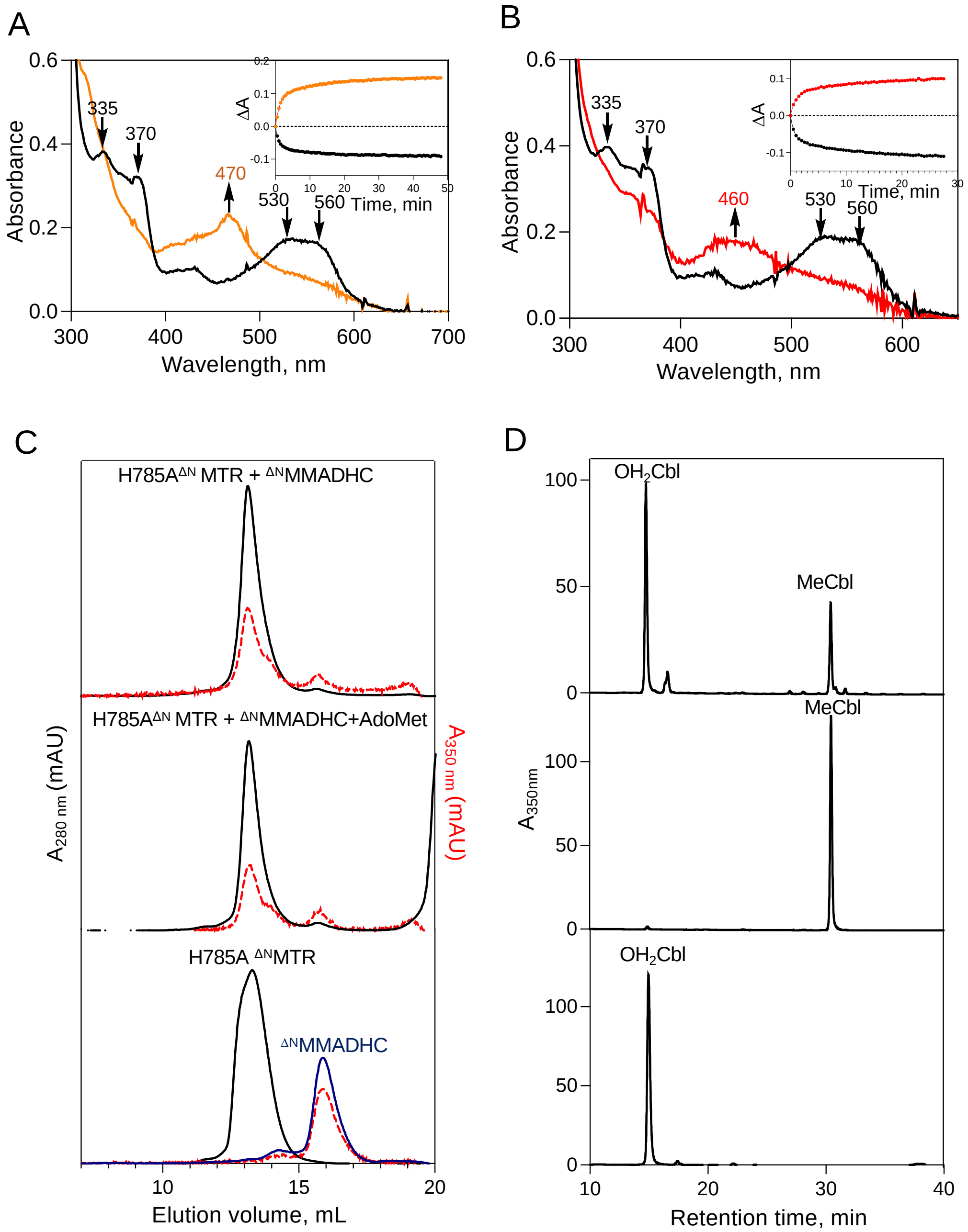


**Figure S4. Cobalamin transfer from MMADHC to H785A MTR . A.** $^{\Delta N}$MMADHC-S-cob(III) (20 μM, black trace) was mixed with H785A $^{\Delta N}$MTR (40 μM) in Buffer A at 20°C. Spectra were recorded every 30 s for 50 min and the final spectrum (orange trace) is shown. *Inset.* The dependence of $A_{470\ nm}$ (orange) and $A_{555\ nm}$ (black) on time was used to estimate $k_{obs}$. **B.** $^{\Delta N}$MMADHC-S-cob(III) (20 μM, black trace) and 500 μM AdoMet were mixed with 40 μM H785A $^{\Delta N}$MTR in Buffer A at 20 °C under anaerobic conditions. Spectra were recorded every 30 s for 30 min and the final spectrum (red trace) is shown. *Inset* The dependence of $A_{480\ nm}$ (red) and $A_{555\ nm}$ (black) on time was used to estimate $k_{obs}$. **C.** The reaction mixtures from A and B were separated on a size exclusion column and the eluate was monitored at the indicated wavelengths. The gel filtration profile confirmed cofactor transfer from $^{\Delta N}$MMADHC to H785A $^{\Delta N}$MTR ± AdoMet . **D**. HPLC analysis of the reaction mixture in B confirmed MeCbl formation.

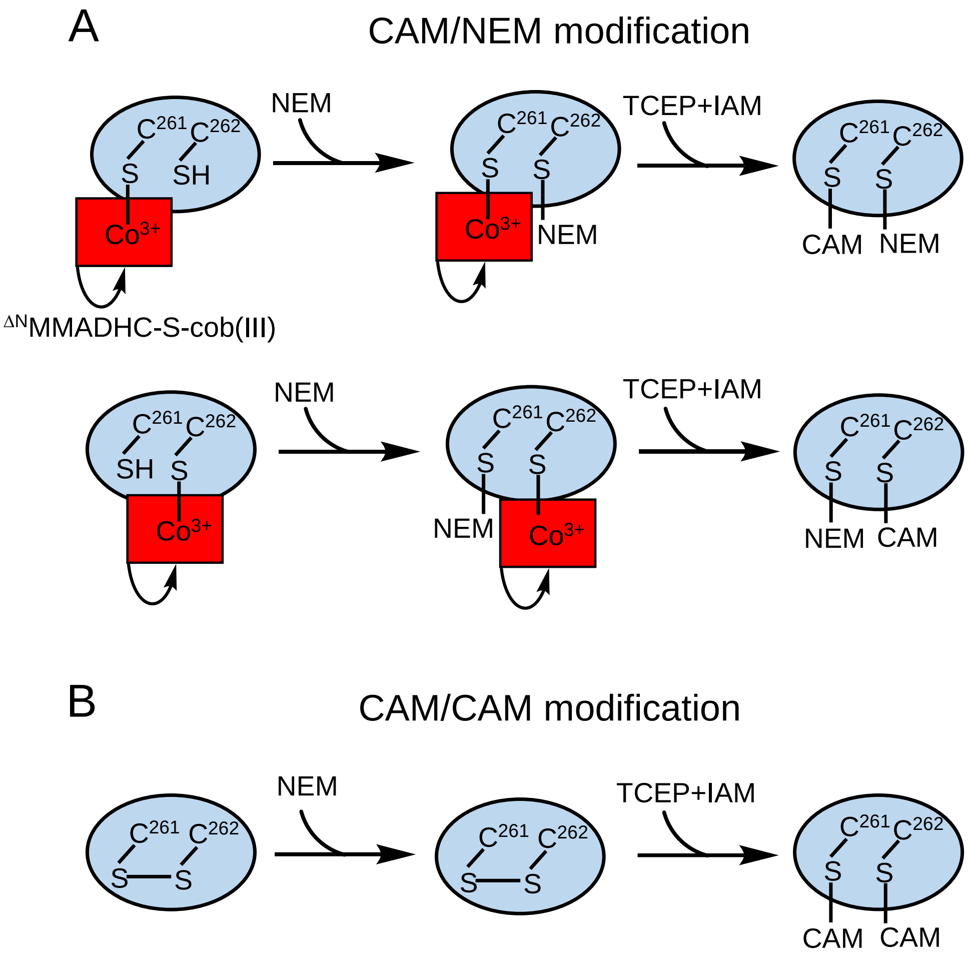


**Figure S5. Thiol labeling strategy for MS analysis**. **A**. Cobalamin can be coordinated to either C261 or Cys-262 on MMADHC and the fraction of $^{\Delta N}$MMADHC-S-cob(III) that did not transfer cobalamin to $^{\Delta N}$MTR, led to CAM/NEM (top) or NEM/CAM (bottom) labeling. **B**. $^{\Delta N}$MMADHC containing a disulfide formed after transfer cobalamin to $^{\Delta N}$MTR, was doubly labeled with CAM following reduction with TCEP and alkylation with iodoacetamide (IAM). NEM is N-ethylmaleimide.

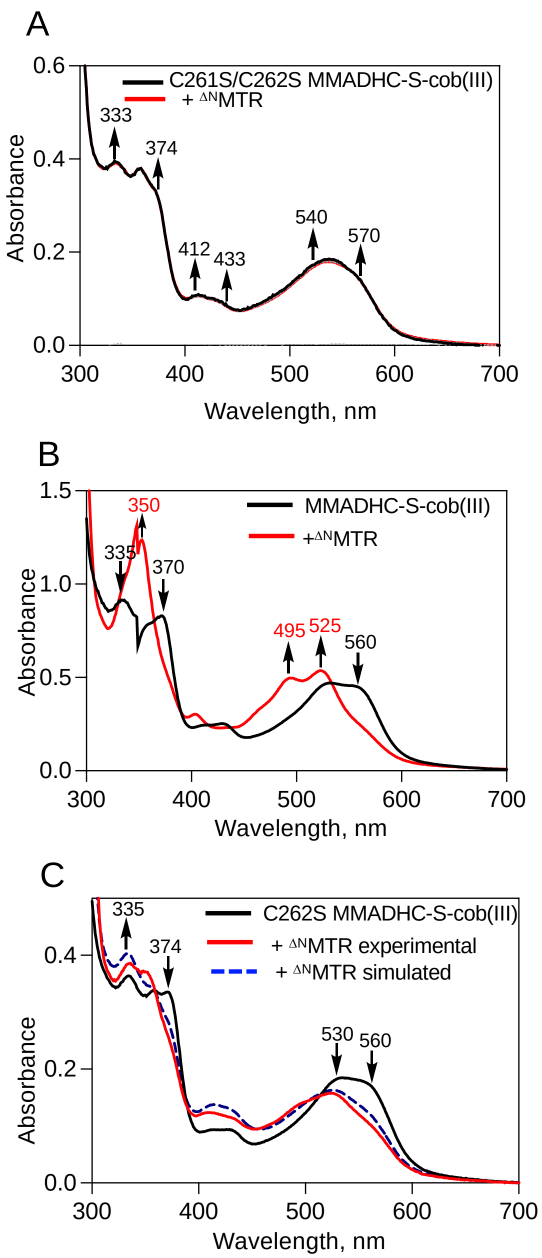


**Figure S6. C261S/C262S MMADHC coordinates $H_2$OCbl. A.** C261S/C262S MMADHC-S-cob(III) (20 μM, black trace) was mixed with $^{\Delta N}$MTR (40 μM) in Buffer A at 20 °C and spectra were recorded every min for 30 min. The final spectrum (red trace) indicates that $B_{12}$ is not transferred from the alternate binding site in the N-terminal disordered region of MMADHC. **B.** MMADHC$^{-}$S-cob(III) (60 μM, black trace) was mixed with $^{\Delta N}$MTR (80 μM) in Buffer A at 20 °C. The final spectrum (red trace) was recorded at 30 min. **C**.C262S MMADHC-S-cob(III) (20 μM, black trace) was mixed with $^{\Delta N}$MTR (40 μM) in Buffer A at 20 °C. The final spectrum (red trace) recorded at 30 min indicates that ~50% of C262S MMADHC$^{-}$S-cob(III) formed a complex with $^{\Delta N}$MTR. A simulated spectrum (blue dashed trace) was calculated by adding the spectra corresponding to 50% MMADHC-S-cob(III) and 50% C262S MMADHC-S-cob(III)-$^{\Delta N}$MTR.
